# Huntingtin polyglutamine expansions misdirect axonal transport by perturbing motor and adaptor recruitment

**DOI:** 10.1101/2024.04.12.589210

**Authors:** Emily N. P. Prowse, Brooke A. Turkalj, Muriel Sébastien, Daniel Beaudet, Heidi M. McBride, Gary J. Brouhard, Mahmoud A. Pouladi, Adam G. Hendricks

## Abstract

Huntington’s disease (HD) is caused by polyglutamine (polyQ) expansions in huntingtin (HTT). Polyglutamine repeat lengths >35Q lead to neurodegeneration and longer repeats correspond to earlier symptom onset. HTT scaffolds kinesin-1 and dynein to a variety of vesicles and organelles directly and through adaptors. To characterize the effects of HTT polyQ expansions on axonal transport, we tracked BDNF vesicles, mitochondria, and lysosomes in neurons induced from an isogenic set of human stem cell lines with repeat lengths of 30, 45, 65, and 81Q. Mild and intermediate pathogenic polyQ expansions caused increased BDNF motility, while HTT-81Q misdirected BDNF towards the distal tip. In comparison, mitochondria and lysosome transport showed mild defects with polyQHTT. We next examined the effect of polyQHTT in combination with neuroinflammatory stress. Under stress, BDNF cargoes in HTT-30Q neurons were more processive. Stress in HTT-81Q resulted in a stark decrease in the number of BDNF cargoes. However, the few remaining BDNF cargoes displayed more frequent long-range motility in both directions. Under neuroinflammatory stress, lysosomes were more abundant in HTT-81Q neurons, and motile lysosomes moved less processively and had an anterograde bias while lysosomes in HTT-30Q where not strongly affected. To examine how HTT-polyQ expansions altered the motors and adaptors on vesicular cargoes, we isolated BDNF cargoes from neurons and quantified the proteins associated with them. BDNF-endosomes isolated from HTT-81Q neurons associated with 2.5 kinesin-1 and 3.9 HAP1 molecules on average, compared to 1.0 kinesin-1 and 1.0 HAP1 molecule for HTT-30Q neurons. Together, these results show that polyQ expansions in HTT cause aberrant motor and adaptor recruitment to cargoes, resulting in dysregulated transport and responses to neuroinflammatory stress.

## Introduction

Huntington’s Disease (HD) is a progressive neurodegenerative disorder caused by polyglutamine expansion mutations at the N-terminal domain of the huntingtin protein (HTT-polyQ)^1^. PolyQ lengths >35Q lead to neurodegeneration, with longer repeats associated with earlier disease onset and increased disease severity^1^. HTT is a scaffolding protein that links cargoes to motor proteins and adaptors. Polyglutamine expansions in HTT result in impaired transport of brain derived neurotrophic factor (BDNF), amyloid precursor protein (APP)-positive vesicles, and autophagosomes^2–4^. Given HTT’s roles in directing the transport of multiple types of intracellular cargoes, we sought to delineate and compare defects in the transport of signalling and degradative organelles.

HTT directs transport by recruiting specific sets of adaptors and interacting proteins to different cargoes. HTT/HTT-associated protein 1 (HAP1) recruit kinesin-1 and dynein to late endosomes, lysosomes, autophagosomes and signalling molecules^5–7^ to drive long-range bidirectional motility. HAP1 interacts with kinesin-1’s C-terminal domain, kinesin-1 adaptor kinesin-light chain (KLC), and dynein adaptor dynactin’s p150^Glued^ subunit^6,8,9^. HTT/HTT-associated protein 40 (HAP40) associates with Ras-related protein 5 (RAB5) to recruit kinesin-1 and -3 and is implicated in switching endocytic and secretory cargoes from microtubule to actin-based transport^10,11^. HTT/optineurin (OPTN) recruits myosin-VI to secretory and golgi-derived cargoes via Ras-related protein 8 (RAB8) to facilitate short-range transport along actin filaments^12,13^. Therefore, polyQ expansions in HTT potentially alter its interactions with multiple adaptors and cargoes.

HTT associates with both signalling and degradative organelles^14,15^. However, each cargo type associates with different motors and exhibits distinct motility characteristics. BDNF vesicles show saltatory, bidirectional movements, driven by kinesin-1, kinesin-3, and dynein^16,17^. HTT/HAP1 mediates dynein activity on retrograde autophagosomes specifically in the mid-axon and is required for their maturation to autolyosomes^15^. The differences in their motility and associated motors suggests that signaling and degradative cargoes are differentially regulated by HTT.

While defects in intracellular transport are widely associated with polyQ mutations in HTT, the direct effects of polyQ expansion length on the recruitment of motors and adaptors is not well characterized. BDNF vesicles, which promote neuronal growth, have decreased velocities, increased pausing, and increased stationary cargoes in neurons with HTT-polyQ^14^. Interestingly, HTT-polyQ has a higher affinity for both HAP1 and the p150^Glued^ subunit of dynactin^14^, suggesting it may sequester or act to inhibit transport through these adaptors. Autophagosomes similarly exhibit decreased velocities, a decreased fraction of cargoes undergoing minus-end transport, and an increase in stationary motility in HD models^4^. Additionally, mitochondria have defective transport or localization in HD^18–21^. Huntingtin knockdown alters the localization of acidified lysosomes^22^, however the effect of HTT-polyQ has not been fully characterized. Here, we combined live cell imaging with experiments on isolated organelles in gene-edited neurons to understand how polyQ length, organelle identity, and adaptor/motor recruitment are altered by polyQ expansions in HTT.

To directly examine the defects in intracellular transport related to HD, we characterized organelle motility and quantified the motors, adaptors, and markers associated with them in neurons differentiated from isogenic H9 human embryonic stem cell (hESC) lines engineered to express endogenous HTT with 30, 45, 65, or 81Q expansions^23^. First, we used immunofluorescence to characterize the localization of HTT and measured the amount of HTT in each cell line. HTT localizes to the cytoplasm at 45Q and 65Q lengths, while its localization in 30Q and 81Q neurons was more punctate. We then tracked the transport of BDNF vesicles, lysosomes, and mitochondria in 30Q, 45Q, 65Q, and 81Q neurons. BDNF cargo transport was affected most strongly, with a shift towards anterograde motility in HTT-81Q neurons. To mimic neuroinflammatory stress, we treated neurons with interferon γ (IFNγ). Upon IFNγ addition, defects in BDNF and lysosome transport with HTT-polyQ were amplified: BDNF cargoes were less abundant, misdirected, and more motile, while lysosomes were more abundant, misdirected, and more frequently stationary. We then characterized BDNF cargoes in vitro and showed that 81Q neurons recruited more kinesin-1 and HAP1 to BDNF vesicles than control 30Q, suggesting differences in their transport complex composition. These experiments further reveal HTT’s regulatory role in axonal transport and delineate how it becomes disrupted in disease.

## Results

### Huntingtin levels and density increase with HTT-polyQ

To examine the effect of polyQ length on neuronal morphology and HTT localization, we characterized neurons by immunofluorescence. We imaged cortical neurons derived from 30, 45, 65, and 81Q HTT isogenic embryonic stem cells 7 days post terminal differention (∼DIV7). At this stage, neurons exhibited a defined axon compared to shorter neurites and had limited synaptic formation. These isogenic lines were previously characterized by the Pouladi lab, where markers for pluripotency (OCT4, LIN28, NANOG, SSEA4 markers), neural progenitor identity (FOXG1, NESTIN, PAX6 markers) and neuronal differentiation (MAP2) indicated no significant differences in pluripotency nor differentiation^23^. However, the 81Q cells had fewer medium spiny neuron marker DARPP32 transcripts by quantitative PCR and by immunofluorescence after differentiation to cortical neurons, indicating that the other cell types contained some striatal neurons^23^. After optimizing cell culture conditions for differentiation including cell media and laminin concentrations, we performed immunofluorescence and morphological characterization of each cell line and noted no differences in differentiation or pluripotency between cell types. We imaged neurons at low magnification for HTT, a neuron-specific isotype of tubulin (β3-tubulin), and DAPI to identify nuclei, and calculated their arborization lengths, HTT levels and localization as a function of polyQ length (**Figure 1A, S1**). The HTT intensity normalized to the tubulin signal increases by 19.6% and 2.36% for 65Q or 81Q HTT, respectively (**Figure 1B**). 45Q-HTT neurons have longer neurites than other neurons (279% above control, **Figure 1C**), consistent with HTT’s roles in development and neurogenesis^24–26^. Next, we compared the amounts of HTT in the mid-axon localized to the cytosol or in punctae, where puctae may indicate vesicle association or HTT aggregation. To quantify HTT localization, we used linescan analysis of higher magnification images (**Figure 1D, E**). We then measured the variance of intensities along the axon, where the variance is driven by the difference in intensities between bright punctae and diffuse cytosolic localization. High intensity variance indicates more punctate localization. Interestingly, the variance and HTT intensity values decrease in the 45Q and 65Q neurons by -51.4% and -49.5% respectively, which suggests HTT’s association with vesicles is weakened and its localization shifts to the cytoplasm (**Figure 1F, G**). In 81Q neurons, the high variance (38.7% above control) and high intensity values likely indicate elevated HTT self-association into aggregates which may be cytosolic or clustered with vesicular cargoes (**Figure 1 F, G**). Overall, immunofluorescence characterization demonstrates that HTT levels increase with extended polyQ lengths ^23^, while its localization shifts to the cytoplasm at intermediate lengths and we propose it aggregates more readily in HTT-81Q neurons.

**Figure 1.**
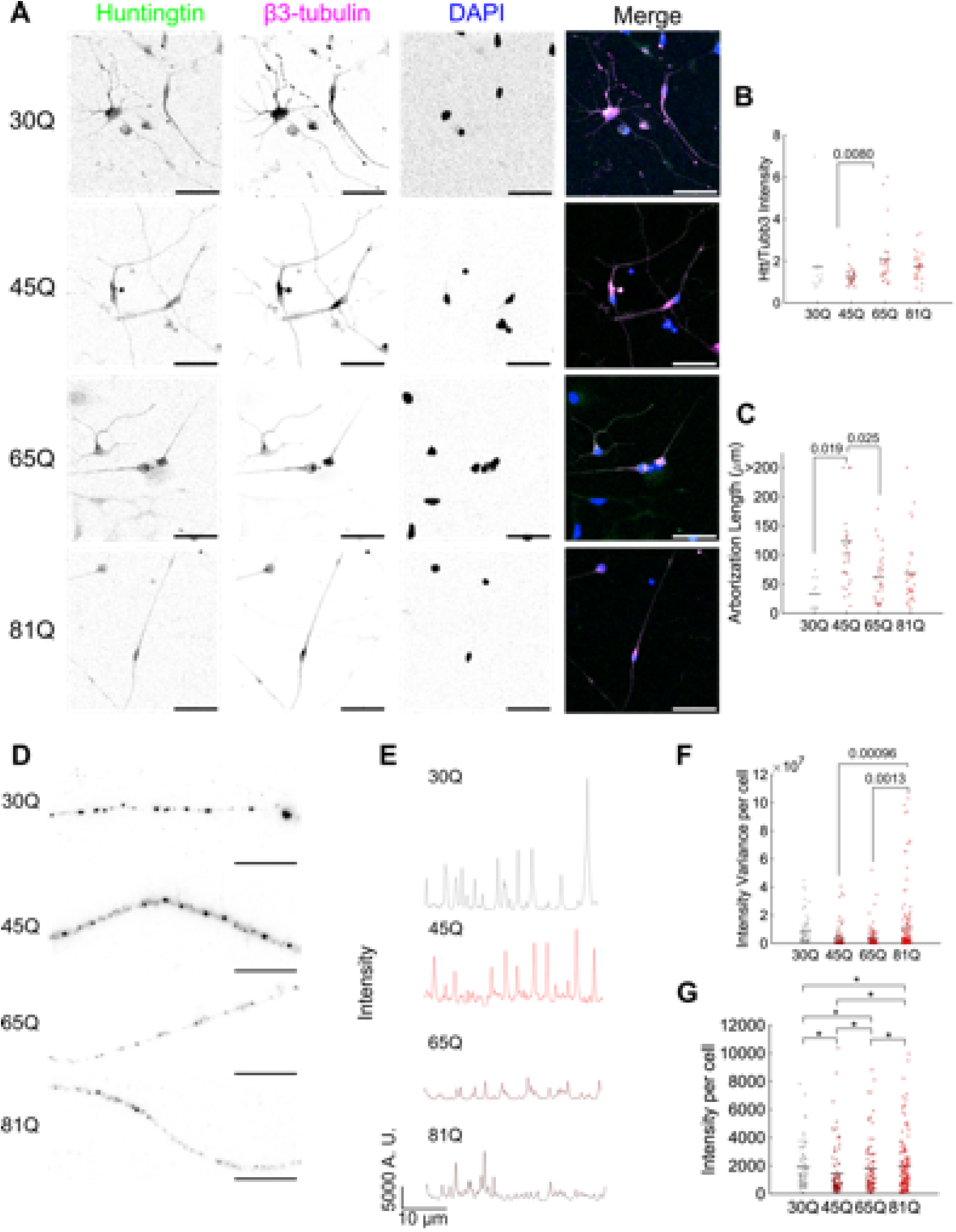
Polyglutamine expansions increase the intensity of HTT punctae. **A** Immunofluorescence images showing HTT (green), β3-tubulin (magenta), DAPI (blue), and merged images for neurons of 45Q (top), 65Q (middle), and 81Q (bottom) cell lines. Scale bar 50 μm. **B** Relative HTT intensity to β3-tubulin quantified from images as in **A** (30Q, n=12 images, 2 terminal differentiations, 45Q n=32 images, 3 terminal differentiations, 65Q n=31 images, 3 terminal differentiations, 81Q n=30 images, 3 terminal differentiations), mean values: 30Q 1.72, 45Q 1.26, 65Q 2.05, and 81Q 1.76. Means are indicated by black horizontal line. **C** Quantification of axon arborization lengths for each cell line above from the same images as in **A** and **B**, mean values: 30Q 32.6, 45Q 124, 65Q 61.4, and 81Q 68.5 μm. **D** Immunofluorescence images of HTT in 30Q, 45Q, 65Q, and 81Q HTT neurons at higher magnification, scale bar 10 μm. **E** Linescan intensity profiles with a linewidth of 10 pixels for each respective image in **D**. **F** Mean variance in intensity per cell in linescan analysis from 30Q (n=72 neurons, 2 terminal differentiations), 45Q (n=90 neurons, 3 terminal differentiations), 65Q (n=90 neurons, 3 terminal differentiations), and 81Q (n=138 axon segments from ∼100 neurons, 2 terminal differentiations). Mean values for intensity variance: 30Q 8.11 ×10^6^, 45Q 3.94 × 10^6^, 65Q 4.09 × 10^6^, and 81Q 11.2×10^6^. **G** Mean intensity per cell for all linescan images, mean values: 30Q 1.91 × 10^3^, 45Q 1.46 × 10^3^, 1.80 × 10^3^, and 1.98 × 10^3^. Statistical significance was determined by analysis of variance (ANOVA) followed by Tukey’s multiple comparison post-hoc test, p values at 95% confidence are listed where possible and otherwise are indicated by an asterisk (*). Values for the asterisks in **G** are all < 0.001.

### Huntingtin polyglutamine expansions alter BDNF vesicle motility and direction

BDNF vesicles are transported bidirectionally along the axon. Striatal neurons are most susceptible in HD. Striatal neurons do not produce BDNF, and instead rely on its efficient transport from cortical neurons as their sole source of this pro-survival factor^27–31^. Cortical neurons synthesize the signaling molecule BDNF in the soma and transport it via a secretory path to axon terminals and synapses with striatal neurons^32,33^. Striatal neuron death in HD is attributed to impaired BDNF transport in cortical neurons^27^. We labelled BDNF cargoes using BDNF-coated quantum dots and observed their motility along the mid-axon of neurons 7 days post terminal differentiation (**Figure 2A**). BDNF cargoes exhibited frequent directional switches in all conditions, however the net displacement slightly increased in 45Q and 65Q-HTT neurons (**Figure 2A, S2B**). We used multiple tools to characterize motility over short and long length scales. TrackMate^34^ (**Figure S2**) provides sub-pixel localization (resolution ∼34 nm for BDNF and lysosomes, ∼84 nm for mitochondria) to assess motility over short length scales. However, TrackMate often failed to accurately link the long trajectories observed in neurons. Thus, in parallel, we analyzed kymographs using KymoButler^35^, which is better able to link long trajectories but is limited to nearest-pixel resolution (160 nm).

**Figure 2.**
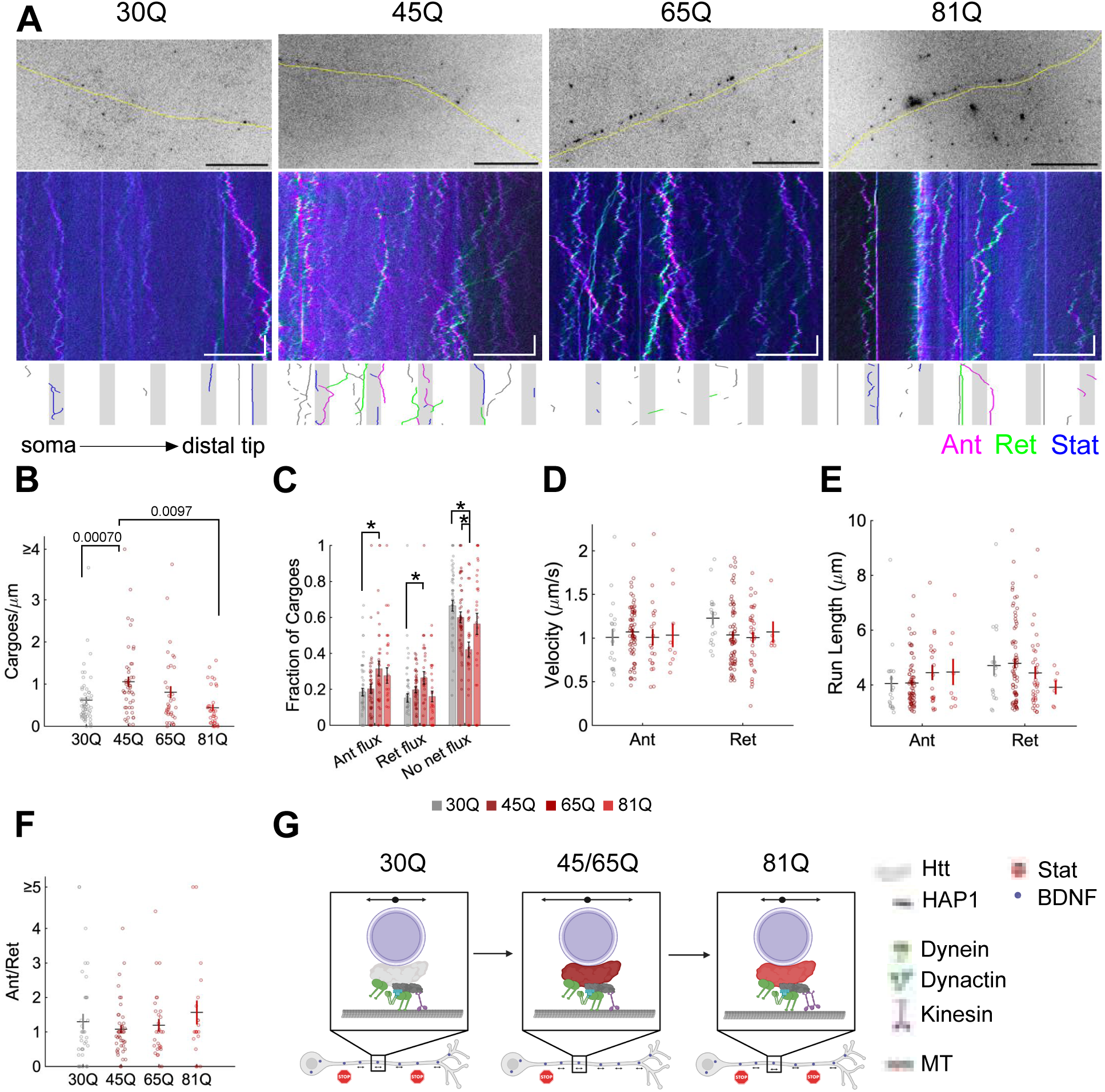
BDNF cargo transport is differentially regulated by intermediate and long pathogenic polyglutamine lengths. **A** Inverted image of the first frame of BDNF-quantum dots in the axon with the shape of the axon traced in yellow below for 30Q, 45Q, 65Q, 81Q neurons (top). Colour coded kymograph generated using KymographClear^69^ demonstrates anterograde (magenta), retrograde (green) and stationary (blue) cargoes for each sample axon image. Scale bars 20 μm, 5 s. Below the kymograph is a plot demonstrating flux characterization of each trajectory continuing in the following frames: gray shaded areas represent the boxes used to calculate flux and the trajectories are coloured according to the flux calculated for the entire trajectory, gray trajectories are not counted because they do not enter the boxes, magenta represents anterograde, green represents retrograde, and blue represents no net flux. **B-F** Long-range track analysis using KymoButler^35^ from 30Q (n=56 neurons, 4 terminal differentiations),45Q (n=47 neurons, 2 terminal differentiations),65Q (n=35 neurons, 2 terminal differentiations), and 81Q (n=34 neurons, 4 terminal differentiations) . Bar heights and horizontal lines indicate the mean of the data, while error bars and vertical lines represent the SEM. **B** Average total number of cargoes per cell when the box width for flux analysis is set to 5 μm (means: 0.615, 1.05, 0.806, and 0.439 for 30, 45, 65, and 81Q, respectively). **C** Fraction of trajectories with anterograde (ant), retrograde (ret), and no net flux resulting in flux averaged per cell, with a 5 μm box. Means for anterograde flux: 30 Q 0.184, 45Q 0.203, 65Q 0.315, 81Q 0.276, retrograde flux: 30Q 0.151, 45Q 0.197, 65Q 0.264, 81Q 0.160, and no net flux: 30Q 0.665, 45Q 0.600, 65Q 0.421, and 81Q 0.564. (**D)** Mean velocity (anterograde: 30Q 1.01, 45Q 1.07, 65Q 1.01, and 81Q 1.03 μm/s, retrograde: 30Q 1.23, 45Q 1.03, 65Q 1.01, and 81Q 1.07 μm/s) and (**E**) run length (anterograde: 30Q 4.04, 45Q 4.07, 65Q 4.45, 81Q 4.47 μm and retrograde: 30Q 4.71, 45Q 4.79, 65Q 4.44, and 81Q 3.91 μm) for each trajectory above 3 μm in run length, separated into runs with a net anterograde (ant) or net retrograde (ret) directionality. Colour code is indicated in the center of the figure, above **G**. **F** Overall directionality by the fraction of anterograde (ant) to retrograde (ret) flux per cell with a 5 μm box width (means: 30Q 1.30, 45Q 1.08, 65Q 1.19, and 81Q 1.56). **G** Model demonstrating the effects on BDNF cargoes in control, intermediate length, and long polyQ HTT. and their length represents processivity or run length of the cargo, vesicles within the axon represent abundance and the arrows or stop signs indicate whether they are motile. Statistical significance with p<0.05 is indicated by the p value itself or an * and was determined by one-way analysis of variance and a Tukey multiple comparison post-hoc test. The values for those with asterisks in **B** are: anterograde flux 30Q vs 65Q, p=0.016, retrograde flux 30Q vs 65Q p=0.0096, For no net flux 30Q vs 65Q p=0.000055, 45Q vs 65Q p=0.0083.

We performed a flux analysis on the KymoButler data by drawing five boxes at different locations along the axon. Trajectories that exited at the somal side were characterized as retrograde, those that exited on the distal side as anterograde, and those that never reached either edge of the box as no net flux (**Figure 2A, bottom panel**). Additionally, cargoes that entered from the soma side but did not exit were counted as anterograde, while cargoes that entered from the distal side and did not exit were counted as retrograde. After comparing box widths from 1-15 µm, we focused our analysis on a box width of 5 μm as the density of cargoes inside the box remained constant above this size and larger boxes resulted in few trajectories traversing the box (**Figure S2D-G**). We observed an increase in the mean number of BDNF vesicles in the 45Q condition, suggesting the longer repeats may have distinct changes to in motor or adaptor composition (**Figure 2B**). All pathogenic polyQ conditions showed increased motility with fewer cargoes exhibiting no net flux (-9.7%, -36.7%, and -15.1% fold changes for 45Q, 65Q, and 81Q, respectively) and less diffusive cargoes (**Figure 2C, S2C**). The fluxes for 45Q and 65Q neurons demonstrated similar trends, with higher directional fluxes compared to controls: 45Q had 10.1% higher anterograde and 30.3% higher retrograde fluxes, while 65Q showed increases by 71.2% for anterograde and 74.4% for retrograde flux (**Figure 2C, Figure S2D-F**). The anterograde flux increased in 81Q (49.9%) neurons as well, however their retrograde flux was lower (only 5.6% above control), creating a 20.7% higher anterograde bias than controls (**Figure 2C, 2F**). The effects of increased uptake in 45Q and elevated motility in both 45Q and 65Q neurons may be a mechanism the neurons use to compensate for the presence of pathogenic HTT. We observed increased long-range transport of BDNF cargoes in both directions for HTT-45Q and HTT-65Q neurons (**Figure 2G**). To generate the observed change in directional bias (**Figure 2C, 2F**), 81Q-HTT may be recruiting or activating more kinesins specifically, perhaps through its association with HAP1 (**Figure 2G**). The exogenous BDNF-endosomes may also be sorted into different endosomal pathway in 30Q-HTT compared to 81Q-HTT neurons. The misdirection of BDNF transport by polyQ-HTT we observe here would be expected to impact neuron survival in HD by perturbing BDNF signalling from cortical to striatal neurons.

### Pathogenic HTT-polyQ causes mild defects in lysosome motility

To examine HTT-polyQ’s impact on degradative cargoes, we tracked lysosomes in the mid-axon of the isogenic HTT-polyQ neurons. Maintaining lysosome transport is essential in neurodegenerative diseases to ensure efficient degradation of cellular waste and protein aggregates, particularly when neurons are under stress. HTT phosphorylation enhances lysosome processivity^36^, and HTT-polyQ has been shown to slow transport of immature lysosomes and autophagosomes^4,37^. We labelled acidified lysosomes with LysoTracker™ and tracked their motility using TrackMate (**Figure S3H-K**). Lysosomes had longer segments of processive motility compared to BDNF vesicles, and were typically biased towards the retrograde direction in control neurons, while we observed more stationary cargoes with longer HTT-polyQ, and an increase in the subpopulation of small lysosomes with fast, processive anterograde runs (**Figure 3A**). We did not observe significant differences in processivity, however displacement increased in the 81Q condition (**Figure S3A, B**). Diffusive lysosome fractions decreased, leading to an increase in stationary fraction for the 45Q condition, while the same trend was observed to a lesser degree as HTT-polyQ length increased (**Figure S3C**). When characterizing long range motility using KymoButler, retrograde velocities in 81Q increased (8.3% from control), while no significant differences in abundance or directionality between cell lines (**Figure 3, S2D-G**). However, the ratio of anterograde to retrograde flux increased in response to HTT-polyQ (72.4%, 140%, and 64.0% above control for 45Q, 65Q, and 81Q, respectively), suggesting HTT-polyQ increases the subpopulation of anterograde-directed lysosomes (**Figure 3F**). In all, the effect of HTT-polyQ on lysosome transport was relatively subtle compared to BDNF endosomes, where lysosomes exhibit a higher fraction of anterograde to retrograde motility in HTT-polyQ neurons (**Figure 3G**).

**Figure 3.**
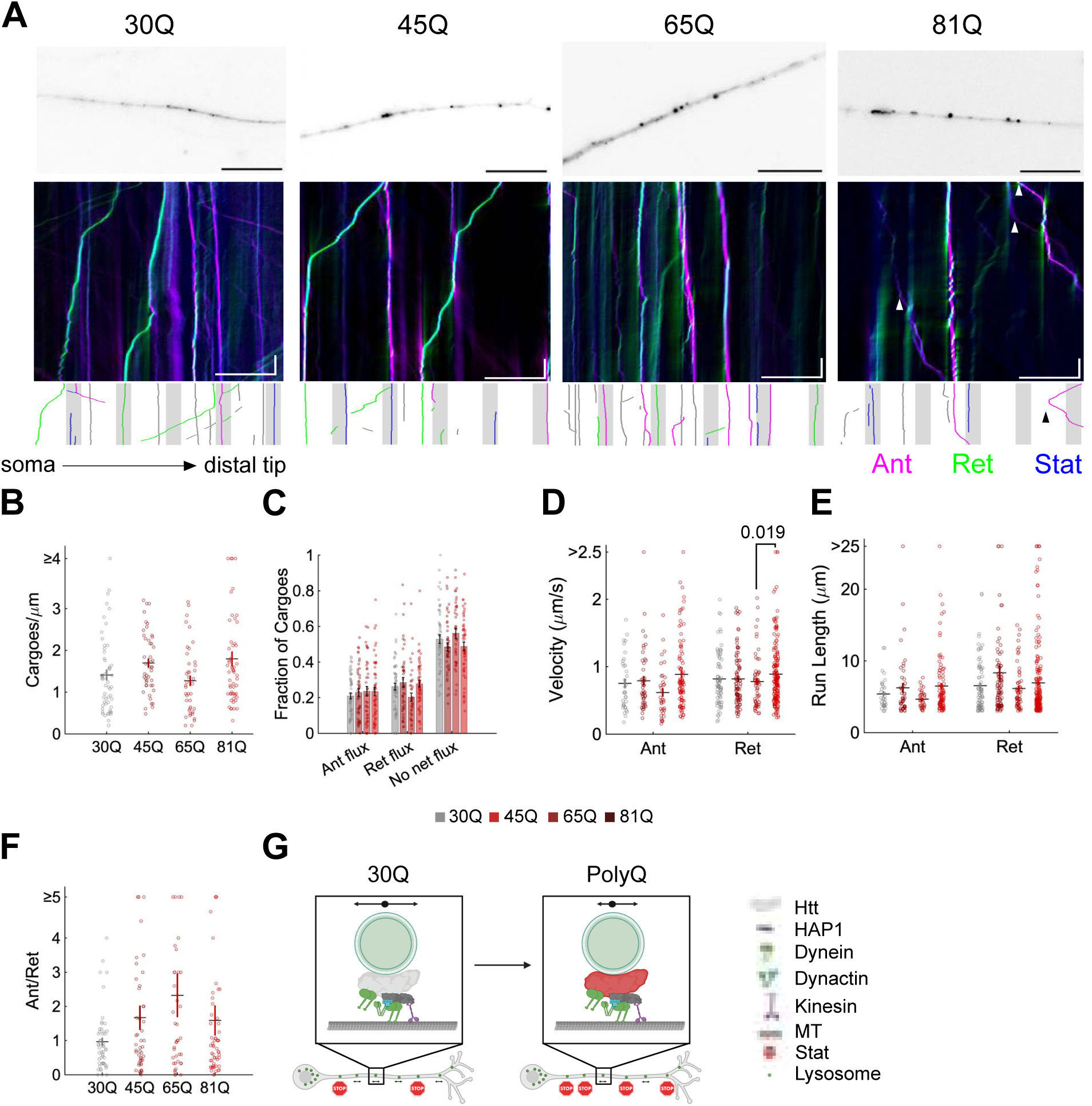
Lysosome transport is mildly misdirected by pathogenic polyglutamine lengths. **A** Inverted images of the first frame of LysoTracker positive lysosomes in the axon for 30Q, 45Q, 65Q, and 81Q neurons (top). Colour coded kymograph generated using KymographClear^69^ demonstrate anterograde (magenta), retrograde (green) and stationary (blue) cargoes for each sample axon image (top). Arrowheads indicate persistent anterograde motility and scale bars are 20 μm, 5 s. Below the kymograph is a plot demonstrating flux characterization of each trajectory continuing in the following frames: gray shaded areas represent the boxes used to calculate flux and the trajectories are coloured according to the flux calculated for the entire trajectory, gray trajectories are not counted because they do not enter the boxes, magenta represents anterograde, green represents retrograde, and blue represents no net flux. **B-F** Long-range track analysis using KymoButler from 30Q (n=54, 3 terminal differentiations),45Q (n=46, 2 terminal differentiations),65Q (n=41, 4 terminal differentiations), and 81Q (n=55, 5 terminal differentiations). Bar heights and horizontal lines indicate the mean of the data, while error bars and vertical lines represent the SEM. **B** Average number of cargoes per micron in each cell when the box width for flux analysis is set to 5 μm (means: 30Q 1.41, 45Q 1.70, 65Q 1.27, and 81Q 1.80). **C** Fraction of trajectories with anterograde (ant), retrograde (ret), and no net flux resulting in flux averaged per cell, with a 5 μm box. Means for anterograde flux: 30Q 0.209, 45Q 0.229, 65Q 0.236, and 81Q 0.235, retrograde flux: 30Q 0.262, 45Q 0.286, 65Q 0.203, and 81Q 0.277, and no net flux: 30Q 0.529, 45Q 0.485, 65Q 0.561, and 81Q 0.488. **D, E** Mean velocity (**D,** anterograde means: 30Q 0.755, 45Q 0.791, 65Q 0.619, and 81Q 0.886 μm/s, retrograde means: 30Q 0.821, 45Q 0.819, 65Q 0.778, and 81Q 0.888 μm/s) and run length (**E,** anterograde means: 30Q 5.41, 45Q 6.24, 65Q 4.67, and 81Q 6.48 μm, retrograde means: 30Q 6.56, 45Q 8.34, 65Q 6.18, and 81Q 6.95 μm) for each trajectory above 3 μm in run length, separated into runs with a net anterograde (ant) or net retrograde (ret) directionality. Colour code is indicated in the center of the figure, above **G**. **F** Overall directionality by the fraction of anterograde (ant) to retrograde (ret) flux per cell with a 5 μm box width (means: 30Q 0.968, 45Q 1.67, 65Q 2.32, and 81Q 1.59). **G** Model demonstrating the effects on lysosomes in control, and long polyQ HTT. and their length represents processivity or run length of the cargo, vesicles within the axon represent abundance and the arrows or stop signs indicate whether they are motile. Bar height or horizontal line indicate the mean of the data, error bars represent the SEM. Statistical significance with p<0.05 is indicated by the p value itself and was determined by one-way analysis of variance and a Tukey multiple comparison post-hoc test.

### Mitochondria motility is largely unaffected by polyQ expansions

We next examined the effect of HTT-polyQ on mitochondrial motility. Changes to mitochondrial morphology and dynamics have been noted in models of HD, although it is unclear whether full-length, non-pathogenic HTT associates directly with mitochondria ^20–23^. We expected the effect on mitochondrial transport to be mild due to the lack of response to S421 phosphorylation which otherwise impacts the directional bias of neuronal HTT cargoes^24^. In addition, Miro/TRAK have been demonstrated as the primary regulators of mitochondrial transport^38–40^. Mitochondria positive for MitoTracker™ were less dynamic than BDNF vesicles or lysosomes, with shorter runs and more stationary cargoes (**Figure 4A**). They were not strongly affected by HTT polyQ expansions (**Figure 4A**). Short-range transport analysis showed slightly decreased processivity and displacement (**Figure S4A-B, H-K**). The processive fraction slightly decreased in all pathogenic HTT-polyQ neurons, suggesting a mild defect in short-range mitochondrial transport (**Figure S4C**). We observed no significant effect compared to 30Q controls on mitochondria number, velocity, run length, flux at 5 μm box width, or directionality from the long-range transport analysis (**Figure 4B-F, S3D-G**). Interestingly, we note the 45Q had various differences in short-range motility compared to the other HTT-polyQ conditions, suggesting HTT-45Q may associate more frequently with mitochondria than the longer HTT-polyQ, which is more subject to self-association/aggregation. The mild defect in processivity observed in response to pathogenic HTT-polyQ expansions (**Figure 4G**) is in close agreement with previous studies^18–21^. From these data, we hypothesize the defects previously observed in mitochondria transport may be indirect effects due to the health of the neuron, while other scaffolds and adaptors including TRAK/Milton act as the primary regulators of mitochondrial transport^38,39,41^.

**Figure 4.**
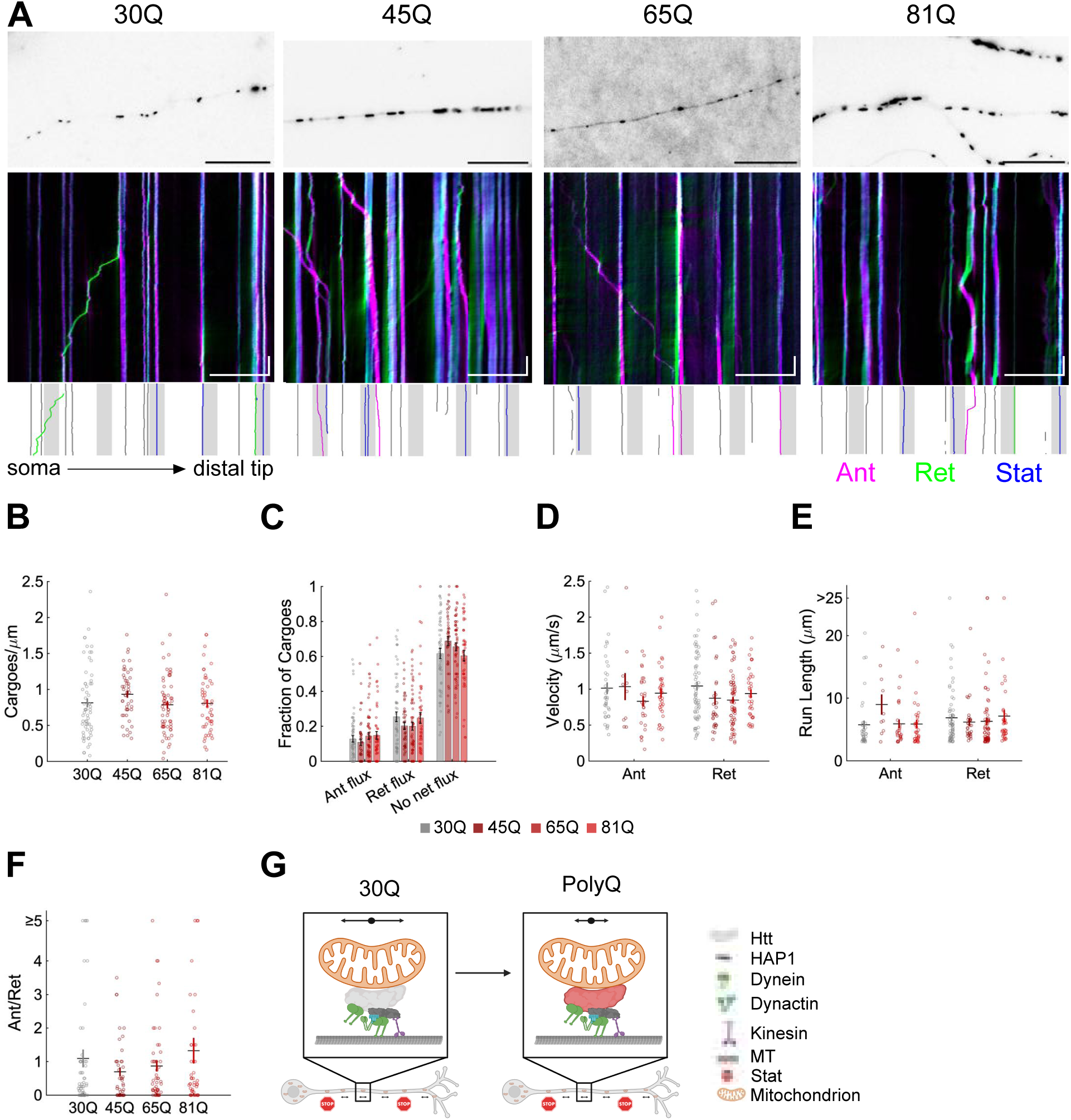
Mitochondria motility is not strongly affected by polyglutamine expansions in HTT. **A** Inverted image of the first frame of MitoTracker positive mitochondria in the axon from HTT with 30Q, 45Q, 65Q, and 81Q neurons (top). Colour coded kymograph generated using KymographClear^69^ demonstrate anterograde (magenta), retrograde (green) and stationary (blue) cargoes for each sample axon image (top). Scale bars 20 μm, 5 s. Below the kymograph is a plot demonstrating flux characterization of each trajectory continuing in the following frames: gray shaded areas represent the boxes used to calculate flux and the trajectories are coloured according to the flux calculated for the entire trajectory, gray trajectories are not counted because they do not enter the boxes, magenta represents anterograde, green represents retrograde, and blue represents no net flux. Scale bars 20 μm, 5 s. **B-F** Long-range track analysis using KymoButler for 30Q (n=61, 5 terminal differentiations), 45Q (n=47, 2 terminal differentiations),65Q (n=64, 3 terminal differentiations), and 81Q (n=50, 5 terminal differentiations) neurons. **B** Average total number of cargoes per micron in each cell when the box width for flux analysis is set to 5 μm (means: 30Q 0.814, 45Q 0.934, 65Q 0.879, and 81Q 0.806) . **C** Fraction of trajectories with anterograde (ant), retrograde (ret), and no net flux resulting in flux averaged per cell, with a 5 μm box. Means for anterograde: 30Q 0.128, 45Q 0.109, 65Q 0.145, and 81Q 0.148, retrograde: 30Q 0.254, 45Q 0.203, 65Q 0.200, and 81Q 0.248, and no net flux: 30Q 0.617, 45Q 0.688, 65Q 0.655, and 81Q 0.694. **D, E** Mean velocity (**D**, anterograde means: 30Q 1.01, 45Q 1.03, 65Q 0.831, and 81Q 0.944 μm/s and retrograde: 30Q 1.04, 45Q 0.975, 65Q 0.848, and 81Q 0.939 μm/s) and run length (**E**, anterograde means: 30Q 5.75, 45Q 8.99, 65Q 5.91, 81Q 5.88 μm, retrograde: 30Q 6.87, 45Q 6.19, 65Q 6.29, and 81Q 7.14 μm) for each trajectory above 3 μm in run length, separated into runs with a net anterograde (ant) or net retrograde (ret) directionality. Colour code is indicated in the center of the figure, above **G**. **F** Overall directionality by the fraction of anterograde (ant) to retrograde (ret) flux per cell with a 5 μm box width (means: 30Q 1.09, 45Q 0.693, 65Q 0.869, and 81Q 1.32). **G** Model demonstrating the effects on mitochondria in control, intermediate length, and long polyQ HTT. and their length represents processivity or run length of the cargo, vesicles within the axon represent abundance and the arrows or stop signs indicate whether they are motile. Bar height or horizontal line indicate the mean of the data, error bars represent the SEM. Statistical significance with p<0.05 is indicated by the p value itself and was determined by one-way analysis of variance and a Tukey multiple comparison post-hoc test.

### Stress in HTT-polyQ neurons alters BDNF and lysosome abundance and direction

A puzzling aspect of many neurodegenerative diseases is that while mutations are often present from birth, symptoms don’t appear until later in life, suggesting a role in accumulated stresses in the development of disease. We used interferon-γ (IFNγ), an inflammatory cytokine, to mimic the neuroinflammation observed in neurodegenerative diseases including HD^42–45^. We exposed the 30Q control or 81Q neurons to 100 ng/mL IFNγ for 24 hours and measured the response in BDNF-quantum dot transport (**Figure 5A**). BDNF vesicles under stress maintained their bidirectional processive motility regardless of polyQ length, however the number of BDNF cargoes increased in 30Q neurons (27.2% higher than control), while it decreased by 58.8% in 81Q neurons (**Figure 5A-B, S5**). This endocytic defect is consistent with previous results showing endocytosis is impaired in HD due to HTT’s interaction with endocytic proteins dynamin 1, HIP1, and HIP1R^46^. Under stress, we expected reduced motility for 81Q neurons and perhaps increased motility in the control neurons, given that BDNF is an anti-apoptotic signal^27^. Interestingly, we observed increased retrograde flux (67.6% above control) upon IFNγ exposure in 81Q neurons and fewer cargoes without net flux compared to the control condition (-27.3%), suggesting BDNF cargoes are more frequently undergoing long range retrograde transport under stress (**Figure 5C**). In characterizing their motility further, we observed mildly increased anterograde velocities (11.25% above control) with shorter run lengths (-13.3%) in 81Q neurons under stress (**Figure 5D, E**). All conditions apart from the control had decreased retrograde velocities (by -18.8%, -12.88%, -13.3% for 30Q+IFNγ, 81Q, 81Q+IFNγ, respectively) and run lengths were reduced by -5.82%, -16.8%, and -13.4% for 30Q+IFNγ, 81Q, and 81Q+IFNγ (**Figure 5D, E**). HTT-polyQ neurons with IFNγ showed increased anterograde compared to retrograde flux (2.83%), suggesting stress maintains the misdirection seen in the 81Q condition despite increasing the fraction of motile cargoes (**Figure 5C, F**). In summary, we observed fewer BDNF vesicles in 81Q neurons under stress, with decreased short range processive transport, but more long-range transport in both directions (**Figure 5G**). Control neurons in the presence of stress showed similar motility characteristics to the 81Q neurons in the absence of stress, suggesting that neurons respond similarly to polyQ expansions and inflammatory stress (**Figure 5B-E**). Taken together, BDNF transport shows opposite responses to neuroinflammation in control and polyQ-HTT neurons, where the number of BDNF cargoes in HTT-30Q neurons increases while the number decreases in HTT-81Q neurons.

**Figure 5.**
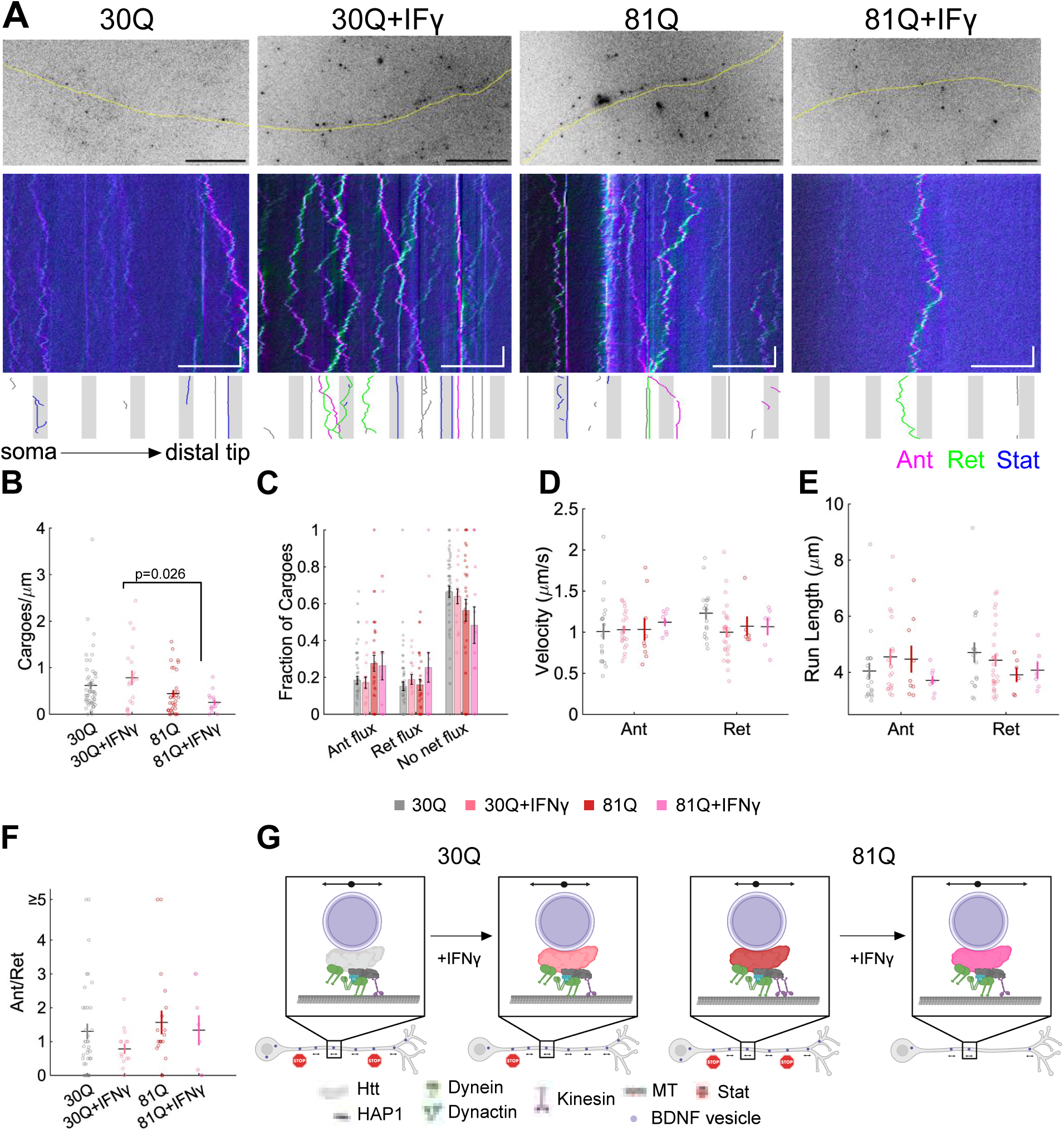
Pathogenic HTT alters the response of BDNF transport to neuroinflammatory stress. **A** Inverted image of the first frame of BDNF-quantum dots in the axon with the shape of the axon traced in yellow below for 30Q-81Q neurons. Colour coded kymograph generated using KymographClear^69^ demonstrate anterograde (magenta), retrograde (green) and stationary (blue) cargoes for each sample axon image (top). Scale bars 20 μm, 5 s. Below the kymograph is a plot demonstrating flux characterization of each trajectory continuing in the following frames: gray shaded areas represent the boxes used to calculate flux and the trajectories are coloured according to the flux calculated for the entire trajectory, gray trajectories are not counted because they do not enter the boxes, magenta represents anterograde, green represents retrograde, and blue represents no net flux. Scale bars 20 μm, 5 s. **B-F** Long-range track analysis for 30Q (same samples as Figure 2, n=56, 4 terminal differentiations), 30Q with 24 hrs 100 ng/mL IFNγ (n=20, 3 terminal differentiations), 81Q (same samples as Figure 2, n=34, 4 terminal differentiations), and 81Q with 24 hrs 100 ng/mL IFNγ (n=15, 3 terminal differentiations) **B** Average total number of cargoes per micron in each cell when the box width for flux analysis is set to 5 μm (means: 30Q 0.615, 30Q+IFNγ 0.782, 81Q 0.439, and 81Q+IFNγ 0.253). **C** Fraction of trajectories with anterograde (ant), retrograde (ret), and no net flux resulting in flux averaged per cell, with a 5 μm box. Bar height or horizontal line indicate the mean of the data, error bars represent the SEM. Mean values for anterograde flux: 30Q 0.184, 30Q+IFNγ 0.171, 81Q 0.276, and 81Q+IFNγ 0.263, retrograde flux: 30Q 0.151, 30Q+IFNγ 0.189, 81Q 0.160, and 81Q+IFNγ 0.254, and no net flux: 30Q 0.664, 30Q+IFNγ 0.640, 81Q 0.564, and 81Q+IFNγ 0.483. **D, E** Mean velocity (**D**, anterograde means: 30Q 1.01, 30Q+IFN 1.03, 81Q 1.03, and 81Q+IFN 1.12 μm/s, retrograde: 30Q 1.23, 30Q+IFNγ 0.999, 81Q 1.07, and 81Q+IFNγ 1.07 μm/s) and run length (**E**, anterograde means: 30Q 4.04, 30Q+IFNγ 4.55, 81Q 4.47, 81Q+IFNγ 3.72 μm, and retrograde means: 30Q 4.71, 30Q+IFNγ 4.43, 81Q 3.91, 81Q+IFNγ 4.08 μm) for each trajectory above 3 μm in run length, separated into runs with a net anterograde (ant) or net retrograde (ret) directionality. Colour code is indicated in the center of the figure, above **G**. **F** Overall directionality by the fraction of anterograde (ant) to retrograde (ret) flux per cell with a 5 μm box width (means: 30Q 1.30, 30Q+IFNγ 0.782, 81Q 1.56, 81Q+IFNγ 1.33). **G** Model demonstrating the effects on BDNF cargoes in control, intermediate length, and long polyQ HTT. and their length represents processivity or run length of the cargo, vesicles within the axon represent abundance and the arrows or stop signs indicate whether they are motile. Statistical significance with p<0.05 is indicated by the p value itself and was determined by one-way analysis of variance and a Tukey multiple comparison post-hoc test.

As with BDNF vesicles, we observed the effects of neuroinflammatory stress on lysosome transport (**Figure 6A**). We expected that cellular stress might upregulate degradative pathways and lysosome transport ^47^. In control neurons, we observed no substantive change in transport under stress (**Figure 6A**). HTT-81Q neurons had an increased number of lysosomes (by 27.3% for 81Q, 53.8% for 81Q+IFNγ compared to controls) and a subpopulation of small lysosomes with persistent anterograde motion regardless of stress (**Figure 6A, B**). Short-range displacements were decreased in 81Q neurons and with IFNγ (**Figure S6A-C**). Long-range lysosome velocities and run lengths were unaffected by stress (**Figure 6D, E**). Lysosome flux is not largely affected by stress in 30Q neurons (**Figure 6C**). However, stress results in reduced retrograde lysosomes in 81Q neurons compared to all other conditions (-22.7% from control), which causes increased anterograde motility (**Figure 6C, F**). Overall, we find lysosome transport is unaffected by stress in control 30Q neurons. In contrast, 81Q neurons responded by increasing lysosome biogenesis and decreasing the fraction of lysosomes being transported towards the soma (**Figure 6G**). Decreased short range transport upon stress appeared universal for both BDNF cargoes and lysosomes, however they differ in the number of these cargoes and their directionality (**Figure 6C, F**, **S6**). Decreased uptake of the pro-survival BDNF and misregulated lysosome biogenesis and transport in 81Q neurons under stress could together contribute to the lower survival of neurons in HD. The anterograde misdirection we observed in both BDNF cargoes and lysosomes expressing HTT-polyQ may be due to increased interactions between HAP1 and HTT^28^ recruiting additional kinesin-1s to cargoes or HTT-polyQ may have a lower affinity for dynein.

**Figure 6.**
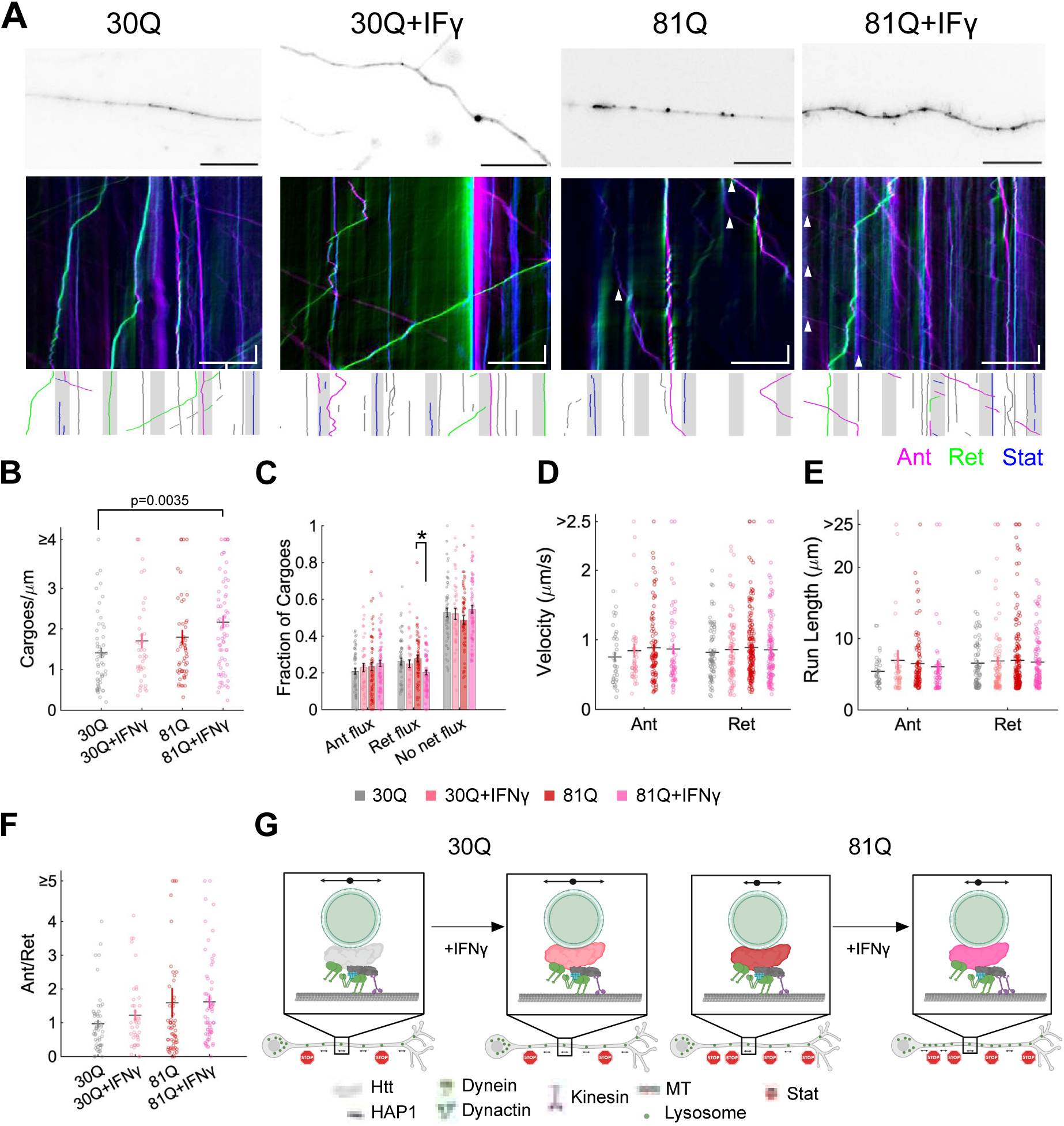
Lysosome biogenesis is upregulated in the presence of both inflammatory stress and pathogenic HTT. **A** First frame image of lysosomes in the mid-axon oriented with their soma on the left and distal tip on the right for 30Q and 81Q neurons in the absence and with 24 hrs treatment with 100 ng/mL IFNγ. Colour coded kymograph generated using KymographClear^69^ demonstrate anterograde (magenta), retrograde (green) and stationary (blue) cargoes for each sample axon image (top). Arrowheads indicate persistent anterograde motility and scale bars are 20 μm, 5 s. Below the kymograph is a plot demonstrating flux characterization of each trajectory continuing in the following frames: gray shaded areas represent the boxes used to calculate flux and the trajectories are coloured according to the flux calculated for the entire trajectory, gray trajectories are not counted because they do not enter the boxes, magenta represents anterograde, green represents retrograde, and blue represents no net flux. **B-F** Long-range track analysis from IFNγ treatment: 30Q samples with IFNγ from n=40 neurons, 2 terminal differentiations, 81Q samples with IFNγ from n=66, 3 terminal differentiations, untreated samples as in Figure 3: 30Q from n=54 neurons, 3 terminal differentiations and 81Q n=55 neurons, 5 terminal differentiations. **B** Average total number of cargoes per micron in each cell when the box width for flux analysis is set to 5 μm (means: 30Q 1.41, 30Q+IFNγ 1.71, 81Q 1.80, and 81Q+IFNγ 2.17). **C** Fraction of trajectories with anterograde (ant), retrograde (ret), and no net flux resulting in flux averaged per cell, with a 5 μm box. Means for anterograde flux: 30Q 0.209, 30Q+IFNγ 0.229, 81Q 0.235, and 81Q+IFNγ 0.251, retrograde flux: 30Q 0.262, 30Q+IFNγ 0.250, 81Q 0.277, 81Q+IFNγ 0.202, and no net flux: 30Q 0.522, 30Q+IFNγ 0.522, 81Q 0.488, and 81Q+IFNγ 0.546. **D, E** Mean velocity (**D**, anterograde means: 30Q 0.755, 30Q+IFNγ 0.843, 81Q 0.886, and 81Q+IFNγ 0.868 μm/s and retrograde means: 30Q 0.821, 30Q+IFNγ 0.859, 81Q 0.888, and 81Q+IFNγ 0.855 μm/s) and run length (**E**, anterograde means: 30Q 5.41, 30Q+IFNγ 6.94, 81Q 6.48, 81Q+IFNγ 6.04 μm, and retrograde: 30Q 6.56, 30Q+IFNγ 6.87, 81Q 6.95, 81Q+IFNγ 6.70 μm) for each trajectory above 3 μm in run length, separated into runs with a net anterograde (ant) or net retrograde (ret) directionality. Colour code is indicated in the center of the figure, above **G**. **F** Overall directionality by the fraction of anterograde (ant) to retrograde (ret) flux per cell with a 5 μm box width (means: 30Q 0.968, 30Q+IFNγ 1.22, 81Q 1.59, 81Q+IFNγ 1.62). **G** Model demonstrating the effects on lysosomes in control, intermediate length, and long polyQ HTT. and their length represents processivity or run length of the cargo, vesicles within the axon represent abundance and the arrows or stop signs indicate whether they are motile. Bar height or horizontal line indicate the mean of the data, error bars represent the SEM. Statistical significance with p<0.05 is indicated by the p value itself or an * and was determined by one-way analysis of variance and a Tukey multiple comparison post-hoc test. The asterisk represents statistical significance for retrograde flux between 81Q vs 81Q+IFNγ with p=0.0068.

### HTT polyglutamine expansions recruit more kinesin-1 and HAP1 to BDNF-endosomes

To examine the effect of polyQ expansions in recruiting adaptors and motors to organelles, we quantified the numbers and types of proteins associated with individual BDNF-endosomes isolated from 30Q and 81Q neurons (Figure 7A). HTT and HAP1 recruit kinesin-1 and dynein to BDNF cargoes^28,48,49^. We employed stepwise photobleaching to determine the number of HTT, HAP1, kinesin-1, and dynein molecules associated with BDNF-endosomes^50–52^. We immunostained BDNF-endosomes for each protein of interest and imaged in epifluorescence until the intensity was indistinguishable from the background. The fluorescence signal from the Alexa 647 labelled secondary antibodies decays over time in a characteristic stepwise manner due to the photobleaching of individual fluorophores. We divided the total number of photobleaching steps by the number of steps for a single molecule, using kinesin-1 (rkin430-GFP) as a standard to estimate the number of proteins associated with isolated vesicles. Our single molecule control from the expression of recombinant kinesin-1 control (sm-kin-1) showed an average of 6.9 distinct steps^52^. The number of HTT molecules bound to BDNF-endosomes isolated from HTT versus HTT-polyQ neurons was similar. However, the number of kinesin-1 motors and HAP1 adaptors associated with 81Q-endosomes was significantly greater than the number associated with 30Q control endosomes, while the number of dynein motors did not change **(Figure 7B)**. (**Figure 7B**). The increased number of kinesin-1 motors is consistent with the increased anterograde motility of BDNF-endosomes we observed in HTT-polyQ neurons (**Figure 2C).** As HAP1 interacts directly with both kinesin-1 and dynein, the observed increase in only kinesin-1 warrants deeper investigation into the mechanism through which polyQ expansions inhibit dynein binding. We then measured the fraction of BDNF-endosomes bound to dynein, kinesin-1, HAP1, and HTT by immunofluorescence colocalization for control and HTT-polyQ endosomes **(Figure 7C)**. The fractions of BDNF-endosome association with dynein, kinesin-1, HAP1 and HTT were normalized to the estimated number of total isolated endosomes for 30Q and 81Q conditions (**Figure S7**, *see methods: **Endosome immunofluorescence and imaging***). A higher fraction of 30Q endosomes associated with dynein, HAP1, and HTT than 81Q endosomes, while 81Q endosomes associated with kinesin-1 more frequently than controls **(Figure 7D)**.

**Figure 7.**
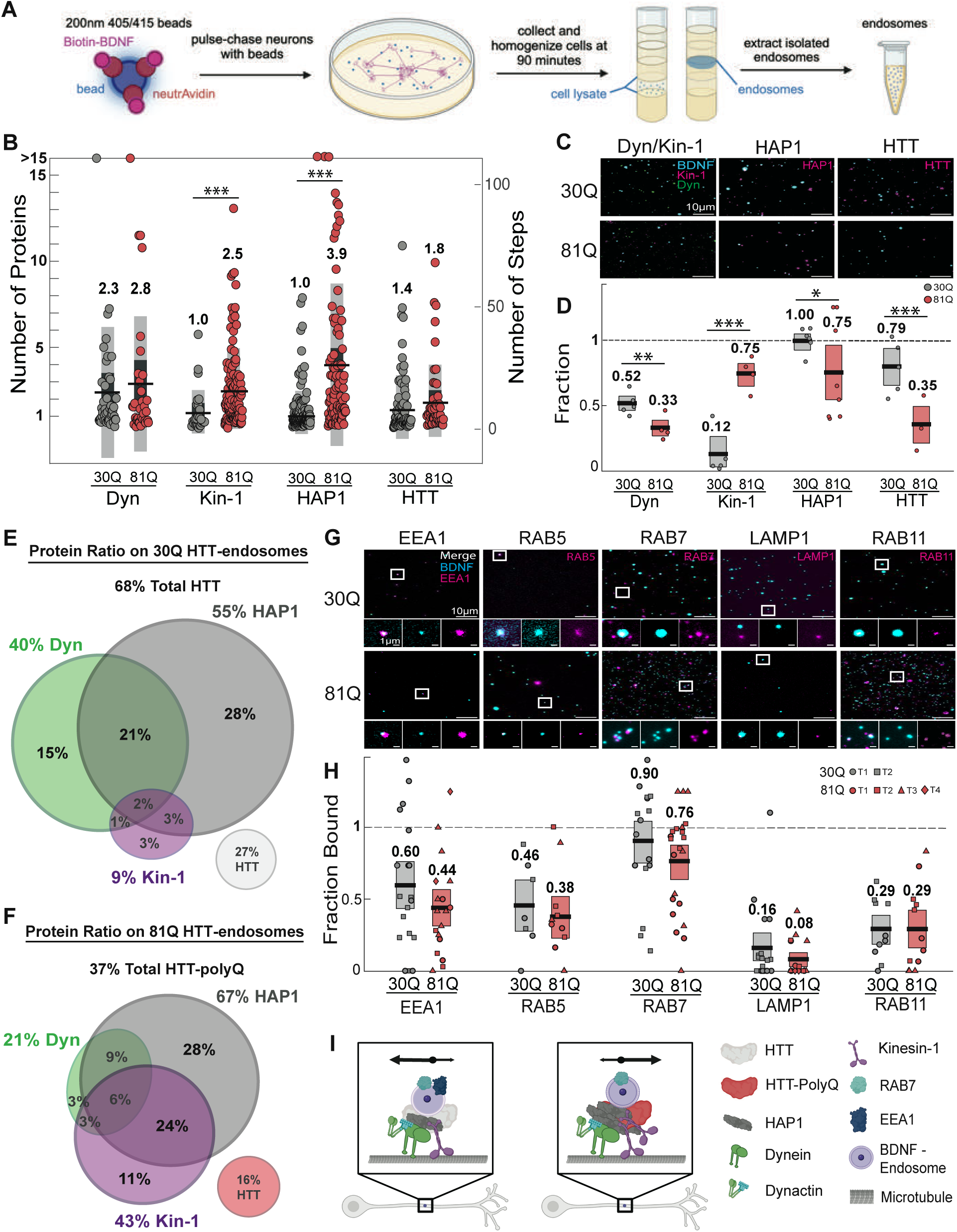
BDNF cargoes in neurons with polyglutamine expanded HTT recruit more HAP1 and kinesin-1 and less HTT. **A** Procedure for isolation of BDNF cargoes, neurons were incubated with beads for 90 minutes. We collected the neurons and homogenized them after the 90 minutes incubation. To obtain the cell lysate, we used sucrose step gradient separation and ultracentrifugation after which isolated endosomes can be extracted^17,25,26,28^. **B** Stepwise photobleaching results from estimations of the number of dynein and kinesin-1 motors, HAP1 adaptors and HTT proteins that associate with 30Q (grey) and 81Q (red) isolated BDNF-endosomes. The plot shows the distribution of the number of dynein, kinesin-1, HAP1 and HTT on 30Q and 81Q isolated BDNF-endosomes. Black lines represent the mean number of proteins, dark grey bars represent SEM values and light grey bars represent 90% confidence intervals. Single-molecule kinesin-1 was used to estimate the number of discrete bleaching steps for a single protein (n=59)^28^. The predictive step count intervals indicate that there were 0-7 dynein on 30Q-endosomes (n=47, 2 isolations) and 0-12 dynein on 81Q-endosomes (n=34, 3 isolations), 0-4 kinesin-1 on 30Q-endosomes (n=27, 2 isolations) and 0-7 kinesin-1 on 81Q-endosomes (n=112, 3 isolations), 0-4 HAP1 on 30Q-endosomes (n=143, 2 isolations) and 0-14 HAP1 on 81Q-endosomes (n=96, 3 isolations), 0-6 HTT on 30Q-endosomes (n=82, 2 isolations) and 0-8 HTT on 81Q-endosomes (n=49, 3 isolations). Bootstrapping of the mean number of steps of each protein associated with 30Q-endosomes compared to 81Q endosomes was performed to test for statistical significance (* p < 0.05, ** p < 0.01, *** p < 0.0001)**. C** Immunofluorescence images show representative fields of isolated BDNF-endosomes (blue) from 30Q (top panel) and 81Q (bottom panel) neurons labelled for dynein (green) and kinesin-1 (magenta), HAP1 (magenta), and HTT (magenta). Scale bars 10 µm. **D** Plot shows fraction of isolated BDNF-endosomes positive for dynein (30Q, n=316 endosomes, 5 fields of view, 2 isolations; 81Q, n=162, 4 fields of view, 2 isolations), kinesin-1 (30Q, n=316, 5 fields of view, 2 isolations; 81Q, n=162, 4 fields of view, 2 isolations), HAP1 (30Q, n=498, 6 fields of view, 2 isolations; 81Q, n=142, 8 fields of view, 2 isolations) and HTT (30Q, n=369, 5 fields of view, 2 isolations; 81Q, n=97, 3 fields of view, 1 isolation) for 30Q (grey) and 81Q (red) neurons. Black lines represent the mean values and boxes show 95% confidence interval for means generated by bootstrapping with replacement. Circles represent the fraction of positive BDNF-endosomes for each field of view. P-values generated from bootstrapped samples were used to test for statistical significance (* p < 0.05, ** p < 0.01, *** p < 0.0001). **E** 30Q and F 81Q bootstrapped, computational model of associations with BDNF-endosomes adjusted for the respective fraction of bead containing endosomes (30Q=68%, 81Q=80%). Venn diagrams show the fractions of HTT-associated endosomes in each population that also exhibit association with dynein (green), kinesin-1 (purple) and/or HAP1 (grey) proteins. The bubbles that do not overlap with the Venn diagram, 30Q (light grey) and 81Q (red), represent the fractions of HTT-associated endosomes that do not associate with any other protein of interest. **G** Immunofluorescence images show representative fields of views with isolated BDNF endosomes (cyan) from 30Q (top panel) and 81Q (bottom panel) neurons labelled with various membrane-associated proteins EEA1, RAB5, RAB7, LAMP1 and RAB11. Below, zoomed in ROIs show merge, BDNF-bead (cyan) and label of interest (magenta). Scale bars 10 µm and 1 µm for selected ROIs. **H** Plot shows fraction of isolated BDNF-endosomes positive for membrane-associated protein stains EEA1, RAB5, RAB7, LAMP1 and RAB11 for 30Q (grey) and 81Q (red) neurons corresponding to intensity plots. Black lines represent the mean values and boxes show 95% confidence interval for means generated by bootstrapping with replacement. 30Q EEA1 (n=566, 2 terminal differentiations), RAB5 (n=79, 2 terminal differentiations), RAB7 (n=460, 2 terminal differentiations), LAMP1 (n=610, 2 terminal differentiations), RAB11 (n=463, 2 terminal differentiations) in grey. 81Q EEA1 (n=293, 4 terminal differentiations), RAB5 (n=346, 3 terminal differentiations), RAB7 (n=365, 2 terminal differentiations), LAMP1 (n=297, 4 terminal differentiations), RAB11 (n=247, 3 terminal differentiations) Shapes represent fraction of positive BDNF-endosomes for each field of view grouped by neuronal terminal differentiation. P-values generated from bootstrapped samples were used to test for statistical significance. **I** Model depicting patterns in association and motility of BDNF-endosomes in cortical neurons expressing 30Q and 81Q HTT. 81Q endosomes associated with more of HAP1 and kinesin-1 compared to 30Q endosomes, increasing anterograde transport.

To account for both the fraction of endosomes associated with a given protein, measured by three-color immunofluorescence, and the average number of proteins on cargoes when the protein is found on the cargo, measured by stepwise photobleaching (*see methods: **Endosome immunofluorescence and imaging***), we used a computational model built from the data in (**Figure 7B-E**) to estimate the ratio of BDNF-endosomes associating with HAP1 (dark grey), kinesin-1 (purple) and dynein (green) in 30Q and 81Q neurons **(Figure 7E, F, S7E)**. An estimated 68% of endosomes in the 30Q population associate with HTT while 37% in the 81Q population associate with polyQ-HTT **(Figure 7E, F)**. The data suggests HTT-polyQ associated endosomes in 81Q neurons preferentially associate with HAP1 and/or kinesin-1, while HTT-associated endosomes in 30Q neurons also associate with HAP1 and/or dynein **(Figure 7E, F)**. The shift in the predominant motor population on endosomes indicates that polyQ expansions in HTT impair the association of HTT to endosomes, and that HTT-polyQ enhances recruitment of kinesins but impairs dynein recruitment to BDNF cargoes (**Figure 7I**).

### BDNF-endosomes associate with early and late endosome markers

Given the changes in transport between 30Q and 81Q BDNF cargoes, we sought to understand if the cargoes were being sorted into different endocytic pathways. Thus, we characterized the markers associated with the endocytosed BNDF-coated beads 90 minutes post-internalization to determine if they were targeted for signaling, recycling, or degradation, and if the endosome maturation pathway differed in 30Q compared to 81Q neurons. We immunostained 30Q- and 81Q-derived BDNF-endosomes with EEA1, RAB5, RAB7, LAMP1 and RAB11 **(Figure 7G)**. ^52,53^. Our results show BDNF-endosome populations primarily associated with late endosome marker RAB7, consistent with previous work in macrophages^51,52^ (**Figure 7H**). Neither 30Q- nor 81Q-endosome populations exhibited strong association with LAMP1, indicating that most isolated BDNF-endosomes were not associated with degradative lysosomes at this timepoint. The intensity distribution means and fraction of association for EEA1, RAB5, RAB7, RAB11, and LAMP1 BDNF-endosomes were not statistically different between 30Q and 81Q populations (**Figure 7H**), indicating BDNF-endosomes are sorted along a similar pathway in both 30Q and 81Q neurons.

## Discussion

Our results provide a direct link between polyQ expansions in HTT and the defects in intracellular transport associated with HD, and highlight HTT’s diverse roles as a scaffolding molecule that links motor proteins and adaptors to multiple organelles. We observe that the effect of polyQ expansions on BDNF transport is dependent on the length of the expansion. Shorter HTT-polyQ may alter motor recruitment or activation on BDNF cargoes, while long 81Q-HTT may be more prone to aggregation which could sequester dynein to misdirect cargo transport. Lysosomes and mitochondria are not strongly affected by HTT-polyQ in normal culture conditions, suggesting other scaffolding complexes may drive their transport. However, under neuroinflammatory stress, HTT-81Q results in defective transport for both BDNF-endosomes and lysosomes. In HTT-81Q neurons under stress, the motility of BDNF cargoes and lysosomes became more anterograde, BDNF cargoes were less frequently endocytosed, and the lysosomes were more abundant. Using quantitative photobleaching, we found that kinesin-1 and HAP1 recruitment were enhanced by HTT-polyQ, consistent with a shift of BDNF cargoes towards the anterograde direction in neurons expressing HTT-polyQ.

While HTT associates with both BDNF and lysosomes, BDNF cargoes exhibited the strongest transport defects associated with HTT-polyQ. Accordingly, we hypothesize that BDNF transport might depend on a more limited set of scaffolding proteins and adaptors compared to lysosomes. In addition, the saltatory and bidirectional motility of BDNF cargoes (**Figure 2A**) might render them more susceptible to changes in motor association or activity compared to the more processive motion of lysosomes or mitochondria in neurons (**Figure 3A, 4A**). Different sets of kinesin motors associate with BDNF cargoes and lysosomes. BDNF cargoes and mitochondria rely on kinesin-1, kinesin-3, and dynein, while lysosomes additionally recruit kinesin-2^16,17,54,55^. Increased anterograde motility of BDNF cargoes in HTT-polyQ neurons (**Figure 2C, F**) is consistent with the more kinesin-1 motors being recruited BDNF-endosomes (**Figure 7B, D**). BDNF-endosome motility is biased towards the retrograde direction, driven by dynein-dynactin^27,28,32,48^. Thus, increasing the fraction of kinesin-1 motors bound to BDNF-endosomes leads to more bidirectional motility. Though HTT associates with lysosomes, lysosomes also associate with other scaffolds and adaptors including RILP/HOOK for dynein-mediated transport and BORC/SKIP/Arl8 for kinesin-mediated transport^56^. The subtle effects we observe on lysosome transport according to HTT polyQ length may be due to a subpopulation that associate and are regulated by HTT, perhaps derived from autophagosomes associated with HAP1. While defects in mitochondria transport have been observed in HD models^18–21^, HTT has not been shown to directly associate with mitochondria. Coupled with the subtle effects of HTT-polyQ on mitochondria transport we observe, these results suggest their transport defects in HD may be indirect. In summary, our results support the model that defective BDNF transport is a primary driver of HD^28^, while the more subtle effects on lysosome transport likely also contribute by impairing degradative pathways.

Previous studies observed defects in BDNF transport upon expression of polyQ-expanded HTT with more frequent pausing, slower velocities, and more stationary cargoes^27–31^. Here, using isogenic stem cell lines edited to express endogenous HTT with varying polyQ expansions, we find that long polyQ expansions in HTT misdirect cargoes anterogradely while shorter polyQ expansions result in enhanced motility compared to controls. These differences may be due to differences in the model systems and/or analysis methods. Multiple previous works used murine cortical neurons with 97-150 polyQ repeats, which correspond with juvenile onset HD and cause severe disease^3,33,57,58^. As we observed, endogenous 81Q-HTT induced changes in transport similar to neuroinflammatory stress while shorter repeat lengths did not, suggesting longer polyQ repeats may cause both direct effects by disrupting normal HTT function and induce cellular stress. Previous studies overexpressed BDNF-mCherry to visualize its transport, which could have altered the number of associated HTTs and motors on each cargo compared to available in the cytoplasm^3,33,57,58^. One study using BDNF conjugated quantum dots in 73Q or 111Q-HTT cell lines showed lower BDNF velocities and lower BDNF abundance in the axon compared to controls via kymograph analysis^59^. Though we observe similar lower abundance of BDNF with HTT-polyQ^59^, particularly under stress, we do not see decreased cargo velocities in either direction. This discrepancy may be due to differences in the analysis, where we performed automated tracking and kymograph analysis on large datasets to examine effects on long-range and short-range motility. Our results show BDNF transport is misdirected with HTT-polyQ compared to previously reported defects resulting in slower or more stationary cargoes.

Huntingtin recruits and regulates motor proteins through adaptors including HAP1, HAP40, optineurin, and HIP1. Our results suggest that HAP1 is required for processive, HTT-mediated BDNF trafficking along microtubules^28,48^. The polyQ repeat region at HTT’s N-terminus is a part of the binding domain for HAP1^49^. PolyQ expansion at the N-terminus as seen in HD may provide additional binding sites for multiple HAP1 adaptors. Previous reports have shown polyQ expansion in HTT leads to an increased interaction between HTT-polyQ with HAP1, and with the dynein-dynactin p150^Glued^ subunit, while association with microtubules was disrupted^28^. In agreement, in 81Q neurons, we observed increased association of BDNF-endosomes with HAP1 and kinesin-1 (**Figure 7B, D**). Additionally, BDNF-endosomes were less frequently bound by 81Q-HTT when compared to BDNF-endosomes in neurons expressing 30Q-HTT (**Figure 7D**), indicating HAP1 binding in excess with HTT-polyQ may hinder additional interactions. The observed increase in kinesin-1 motors over dynein-dynactin may be a result of steric hindrance of HAP1 in the HTT-polyQ condition by the larger 2.4 MDa dynein-dynactin complex versus the smaller 185 kDa kinesin-1^8,51,60^.

In addition to altering the numbers of motors and adaptors recruited by HTT (**Figure 7B**), polyQ expansions also impair the ability of HTT to associate with vesicular cargoes (**Figure 7C,D**). Misfolding of HTT caused by polyQ expansions may result in sequestering of HAP1 and motor proteins and impair interactions with vesicular membranes^61^. Numerous studies of HAP1 overexpression in vitro have reported increased generation of cytoplasmic inclusions, and a tendency for HAP1 to form aggregates and inclusion bodies in physiological contexts^62–64^.

Based on our results, we propose HTT-polyQ misregulates the recruitment of motors and adaptors to organelles and the response to cellular stresses, leading to broad defects in transport of both BDNF and lysosomes (**Figure 5, 6**). Further, HTT-polyQ may induce cellular stress; control neurons under stress have similar BDNF and lysosome transport to 81Q neurons in the absence of stress. When the HTT-polyQ neurons experience inflammatory stress, we observe further defects in both BDNF transport and lysosome transport. With 81Q, stress decreases BDNF endocytosis however increases their anterograde bias and overall motility, while lysosomes are more anterograde and increase in number. The defects in endocytosis and BDNF transport have been previously reported^46^, however we show that BDNF’s increased anterograde motility is driven by association of HAP1 and kinesin-1 with HTT-polyQ and it becomes more pronounced under stress. Given that few BDNF cargoes enter the axon, the increased motility of these cargoes compared to controls under stress could be an attempt to increase BDNF transport efficiency. However, the maintenance of their anterograde directional bias demonstrates that stress cannot rectify the changes to the transport complex induced by HTT-polyQ. We also observe an anterograde bias in lysosome transport with HTT-polyQ under stress and contrary to BDNF, increased lysosome biogenesis. Increasing lysosome biogenesis in response to stress may be a neuroprotective mechanism to prevent infection and reduce misfolded protein populations. In addition, previous works show an increased number of empty autophagosomes in HD^37^, which could increase lysosome demand and supports our claim that HTT-polyQ itself induces a stress response. Frequent anterograde transport of lysosomes in the axon suggests the degradative response is being upregulated and indicates the lysosome-rich population in the soma mobilizes to mitigate the increased degradative demand. HTT-polyQ may activate the stress response and additional stresses cause significant defects in transport of both BDNF and lysosomes.

Given that HTT-polyQ leads to subtle defective responses to inflammatory stress, we suggest that accumulation of minor defects over time lead to significant impairment of neuronal function in HD. If we consider the width of the boxes in our flux measurements, we have a 25 μm segment (5 boxes x 5 μm) that we observe over the course of 3 minutes (**Figures S5D-G & S6D-G**). If we extrapolate the values from **Figure S6E-G** out to one year of a cortical axon of approximately 1 mm in length, this corresponds to a change of 8.4 million more anterograde lysosomes, 12.6 million more without a net flux, and no change in retrograde flux when comparing control (HTT-30Q) to HTT-81Q neurons under stress. For BDNF cargoes, we would have 1.9 million fewer retrograde, 4.2 million fewer without a net flux, and no change to anterograde flux. With these rough estimations, it becomes clearer that the effect of impaired transport can cause surpluses or deficits on the order of millions in the number of cargoes that become misdirected and reach the distal tip or soma over the course of a year, which is tightly controlled in neurons to maintain cell health.

## Materials and Methods

### Cell Culture

The endogenous huntingtin (HTT) gene (27Q) was previously edited to heterozygously express HTT with 30, 45, 65, and 81 polyQ repeats in H9 human embryonic stem cells^23^. We cultured hESCs in mTeSR Plus media (STEMCELL Technologies, Vancouver, BC) on Matrigel (Corning, Corning, NY) coated cell culture dishes using Gentle Cell Dissociation Reagent (STEMCELL Technologies) for passaging. Once confluent, we dissociated with Accutase (Millipore Sigma) induced the stem cells into neural progenitor cells using Stemdiff SMADi induction media (STEMCELL Technologies) within a monolayer over the course of 21-30 days on 10 μg/mL poly-L-ornithine (P3655 Millipore Sigma) in phosphate buffered saline (PBS, Wisent, Saint-Jean Baptiste, QC) and 5 μg/mL laminin (L2020 Millipore Sigma) laminin in Dulbecco’s modified Eagle Medium/ Nutrient Mixture F-12 (Thermo Fisher Scientific) coated-dishes. We cultured neural progenitors in Stemdiff Neural Progenitor Maintenance media (STEMCELL Technologies) or NPC maintenance: 1X N2 (100X, Thermo Fisher Scientific, Waltham, MA), 1X B27 (50X, Thermo Fisher Scientific), 1% MEM non-essential amino acids (Millipore Sigma), 10 ng/mL epidermal growth factor (ab9697, Abcam, Cambridge, MA) 10 ng/mL fibroblast growth factor (10018B Preprotech, Cranbury, NJ), 1 % Antibiotic-antimycotic(Thermo Fisher Scientific) in DMEM/F12 (Thermo Fisher Scientific) for a maximum of 5 passages. We identified no morphological differences between the two media; however, we could maintain the NPCs longer in the Stemdiff Neural Progenitor Maintenance media. When passaging NPCs, we terminally differentiated using BrainPhys Terminal Differentiation media (STEMCELL Technologies) for 6-9 days to generate mature neurons with a clear axon.

### Live Cell Organelle Imaging

We prepared poly-L-orinithine 10 μg/mL in PBS and 5 μg/mL laminin in DMEM/F12 coated coverslips and seeded neural progenitor cells onto them at a low density, i.e., 100,000 per 30 mm round bottom dish, 1 million per 100 mm tissue culture dish containing multiple sterile 25 mm round coverslips. Within the following 0-4 days, we changed the media to BrainPhys (#number STEMCELL Technologies) terminal differentiation media and followed the suppliers’ protocol for further culture. We used neurons for imaging experiments after 6-9 days in terminal differentiation media to obtain clear axonal specification. For stress treatments, we added 100 ng/mL IFγ (ab9659, Abcam) to cell culture for 24 hrs prior to imaging and during imaging. To image lysosomes, we incubated neurons with 70 nM LysoTracker (Thermo Fisher Scientific, Waltham, MA) for 10 minutes prior to replacing the media with BrainPhys with 15 mM HEPES (H3375, Millipore Sigma) as a carbon dioxide buffer for imaging, we then began imaging. We imaged MitoTracker (Thermo Fisher Scientific) by performing the same protocol as with LysoTracker, however with a 20-minute incubation and a concentration of 50 nM during the incubation. For BDNF labelling, we used BDNF-coated quantum dots. We incubated 1 μL of streptavidin-Qdot 565 quantum dots (Q10133MP, Thermo Fisher Scientific) with 10 μL of 10 μg/mL Biotin-BDNF (Alamone Labs, Jerusalem, Israel) and 39 μL of the quantum dot incubation buffer (Q20001, Thermo Fisher Scientific) for a minimum of 30 minutes in the dark on ice. We added 5 μL of the coated quantum dot solution to the pre-existing media on the coverslip and incubated for 30 minutes at 37 C, 5% CO_2_. Next, we removed excess quantum dots in solution by replacing the media with BrainPhys Complete with 15 mM HEPES and began imaging. We imaged the mid-axon inside the Chamlide chamber (Live Cell Instrument, Namyangju-si, Gyeonggi-do, South Korea), defined as >20 μm from the soma or distal tip of the axon, in near-TIRF with a Nikon Ti-E microscope, 100x 1.49 numerical aperture oil immersion objective using 1 mW laser power with 120 ms exposure time on the EMCCD camera for a total of 3 minutes. All samples were imaged within 90 minutes of media replacement with the HEPES buffer.

### Immunofluorescence

We seeded and differentiated neurons on coverslips as described in live cell imaging. After rinsing coverslips with 37°C PBS, we fixed mature neurons using 37°C 4% paraformaldehyde (w/v, Millipore Sigma) diluted in PBS for 8-10 minutes. We next briefly rinsed the coverslips with room temperature PBS and subsequently began incubating with blocking buffer, 2% BSA, 0.2% saponin (Millipore Sigma) in PBS for 30 mins on an orbital shaker at 40-60 rpm. We then incubated the coverslips with the HTT primary antibody (**Table 1**), diluted in blocking buffer at 4°C overnight in a humid chamber. The following day, we washed the coverslips thrice with 2% BSA in PBS for 5 minutes on an orbital shaker at 40-60 rpm. We next incubated the coverslips in the dark with secondary antibody goat anti-mouse 647 in blocking buffer for 45 minutes (dilutions and identifiers in **Table 1**). Next, we washed the coverslips 3 times with PBS protected from light on an orbital shaker at 40-60 rpm for each wash. Finally, for linescan images, we mounted the coverslips on the slide with PBS and began imaging. We imaged the linescan images using the 100X TIRF objective on the DeltaVision OMX in TIRF, with 1.516 refractive index oil, 600 ms exposure time and 2% 647 laser power. For morphological characterization from lower magnification images, we added Tubb3-A647 antibody (**Table 1**) to the secondary antibody mixture, and we used goat-anti-mouse 488 for HTT staining (**Table 1**). Following secondary anitbody incubation, we stained for DAPI (concentrations in **Table 1**) for 20 minutes using a 1μg/ml solution in PBS and rinsed once again with PBS prior to imaging. For the images of the entire axon and multiple neurons for morphological and expression analysis, we mounted the coverslips with ProLong Diamond Antifade Mountant and stored them at 4°C prior to imaging. We imaged these samples using a confocal Zeiss LSM710, with a 10x EC plan neofluoar, NA=0.30, 405 nm blue diode laser 30 mQ, argon ion laser 458/488/514 nm 25 mW, and HeNe red 633 nm 5mW. We also imaged using a 63x 1.4 NA objective.

**Table 1.** Immunofluorescence antibody and stain dilutions with supppliers, identifiers, and concentrations diluted from stocks. When two dilutions are listed, the dilution with the asterisk was used for isolated endosomes and the other dilution was used on fixed cells.

| Reagent | Source | Species | Type | Identifier | Dilution |
| --- | --- | --- | --- | --- | --- |
| Anti-Huntingtin | Abcam | Mouse | Monoclonal | MAB2166 | 1/400, 1/500* |
| Anti-Mouse 647 | Millipore-Sigma | Goat | Polyclonal | A21236 | 1/400, 1/1000* |
| Anti-Tubulin | Millipore-Sigma | Mouse | Monoclonal | T9026 | 1/400 |
| Anti-Tubulin β-3 647 | Biolegend | Mouse | Monoclonal | 657406 | 1/400 |
| Anti-Mouse 488 | Millipore-Sigma | Goat | Polyclonal | A11001 | 1/400, 1/1000* |
| DAPI | ThermoFisher Scientific | N/A | | D1306 | 1 μg/ml |
| Anti-Kinesin-1 | EMD Millipore | Mouse | Monoclonal | MAB1614 | 1/500 |
| Anti-Dynein | Santa Cruz | Rabbit | Polyclonal | sc-9115 | 1/500 |
| Anti-EEA1 | Abcam | Rabbit | Polyclonal | ab109110 | 1/500 |
| Anti-LAMP1 | Invitrogen | Mouse | Monoclonal | MA1-164 | 1/500 |
| Anti-RAB5 | Abcam | Rabbit | Polyclonal | ab18211 | 1/500 |
| Anti-RAB7 | Abcam | Mouse | Monoclonal | ab50533 | 1/500 |
| Anti-RAB11A | Abclonal | Rabbit | Monoclonal | A3251 | 1/500 |
| Anti-HAP1 | Invitrogen | Mouse | Monoclonal | MA1-46412 | 1/500 |
| Anti-Rabbit 488 | Millipore-Sigma | Goat | Polyclonal | 18772 | 1/1000 |
| Anti-Dynein | EMD Millipore | Mouse | Monoclonal | MAB1618 | 1/500 |

### NeutrAvidin Coating of Fluorescent Beads and Validation

We sonicated 250μL of 0.2μm, 365/415nm Fluorospheres_™_ Carboxylate-Modified Microspheres (F8805, Thermo Fisher Scientific) for 5 minutes. Next, we centrifuged the beads for 10 minutes at 14 000 rpm at room temperature and resuspended them in 60 μL of 1.57M EDAC (Bio Basic, Markham, ON), 2.61 M NHS (Bio Basic) in 20 mM 2-ethanesulfonic acid (MES, Bio Basic) with 100 mM NaCl (BioShop, Burlington, ON). We then mixed the resuspended solution using an orbital shaker at 200 rpm, for 3 minutes at room temperature. To wash the beads, we centrifuged them at 14 000 rpm for 2 minutes and resuspended them in 60 μL 200 mM MES with 100 mM NaCl 3 times. After the final wash, we resuspended the beads in 300 μL of 17 μM neutravidin (31000, Thermo Fisher Scientific), 0.001 mg/mL bovine serum albumin (BSA, BioShop), in 0.1 M sodium phosphate buffer at pH=7.0 (Millipore Sigma). We subsequently mixed the solution on an orbital shaker at 200 rpm for 2 hrs at room temperature. We then centrifuged the solution as previously described and washed 3 times by resuspending in 300 μL of 0.001 mg/mL BSA in 0.1 M sodium phosphate buffer, pH 7.0 after each centrifugation. Finally, we centrifuged the bead solution as described previously and resuspended the beads in 200 μL of a filter sterilized solution of cold 0.001 mg/mL BSA in 0.1 M sodium phosphate buffer, pH 7.0. We stored Neutravidin-beads at 4°C for 1-2 months.

### BDNF Conjugation to NeutrAvidin-coated Fluorescent Beads

On the intended day of use, we mixed 20 μL neutravidin-beads with 500 μL BrainPhys terminal differentiation media (STEMCELL Technologies) and sonicated for 5 minutes. 10 μL of 10 μg/mL Biotin-BDNF (B-250-B, Alomone Labs, Jerusalem, Israel), and 3 μL of 10 μg/mL biotin-4-fluorescein (Biotium, Fremont, CA) were added to the solution at a molar ratio ∼1:2-2.5 M and incubated for 30 minutes on ice.

### Endosome Isolation

We plated neurons in coated-cell culture dishes and grew them to ∼80% confluency at 37°C with 5% CO_2_. We performed the endosome isolation as described previously^52,65,66^. We prepared BDNF-beads as described above with BrainPhys terminal differentiation media (STEMCELL Technologies) and incubated it with neurons for 15 minutes. After 15 minutes, we replaced the media to remove beads that have not been internalized, and we incubated the neurons for an additional 75 minutes to isolate endosomes at a late timepoint in transport.

Following bead incubation, we collected neurons using 1 mg/mL Accutase,and spun at 1200 rpm at 4°C for 5 minutes. We resuspended neurons in motility assay buffer (MAB; 10 mM PIPES, 50 mM K-Acetate, 4 mM MgCl_2_, and 1 mM EGTA, pH 7.0.) supplemented with protease inhibitor cocktail (PIC; BioShop), 10mM DTT, 1mM MgATP and 8.5% sucrose. The cell suspension was transferred to a tight-fitting dounce cell homogenizer and homogenized on ice with 50 vertical strokes to ensure adequate disruption of neuronal membranes. Following, we centrifuged the cell homogenate at 2 000 rpm and mixed ∼1 mL of supernatant with 62% sucrose in MAB at a 1:1.2 ratio. All sucrose solutions were supplemented with PIC, 10mM DTT, and 1mM MgATP. We loaded 2 mL of the homogenate mixture on top of a 3.92 ml 62% sucrose in MAB cushion in an Ultra-Clear 5/8” X 4” Beckman Coulter tube (344061, Beckman Coulter). We added the remaining sucrose in MAB solutions: 2.52mL 30% for 200nm beads, 2.61mL 25%, and 2.61mL 10%, on top of the homogenate mixture. Next, we centrifuged the sucrose gradient in a swinging bucket centrifuge with a SWti32 rotor (369650, Beckman Coulter) at 24,000*g* for 72 mins at 4°C. The bead-containing endosomes appeared as a thin band at the 25%–30% sucrose interface and we extracted them using a 21-gauge, BD 305165 PrecisionGlide needle and syringe (14-826C, Fisher Scientific). We kept isolated endosomes on ice at 4°C for up to 24 hours and subsequently stored them at -80°C.

### Endosome immunofluorescence and imaging

We made flow chambers by mounting silanized coverslips to microscope slides using vacuum grease and double-sided tape. Isolated endosomes were added to the flow chambers and incubated for 1 hour at room temperature to allow endosomes to adhere to the surface^67^. Following the incubation, we washed chambers 2X with MAB before and after treatment with Pluronic F-127 (P2443, Sigma-Adlrich) to prevent non-specific binding before incubation with primary antibodies. We incubated endosomes with the combination of primary antibodies for 1) mouse kinesin-1 and rabbit polyclonal dynein, 2) rabbit EEA1 and mouse LAMP1, 3) Rabbit RAB5 and MouseRAB7, 4) Rabbit RAB11A and mouse RAB7, or 5) mouse and 6) mouse huntingtin as identified in **Table 1**, for 1 hour at room temperature in a dark humidity chamber. Following treatment with primary antibodies, we washed endosomes 3X with MAB and incubated with secondary antibodies anti-rabbit Atto 488 and anti-mouse secondary antibody goat anti-mouse Alexa 647 for 1 hour at RT in a dark humidity chamber (Dilutions in **Table 1**). We washed samples 3X with MAB before imaging. To acquire multichannel images, we used brightfield or TIRF with 300 ms exposures using the 405 nm set at 3 mW, and 488 nm and 640 nm lasers set at 1 mW. We selected ROIs based on the centroid values of beads from the parent images captured with 405 nm laser excitation, thresholded according to maximum intensity profiles and scaled to produce circular ROIs with uniform diameters. To choose diameter values, we used the average area in which intensity signal (a.u.) was observed for a sample population of beads. We transferred the ROIs from the parent-bead image to the corresponding images acquired with 565 nm laser excitation. We measured the area and integrated density and saved them in a .csv file. We measured the local background intensity for each ROI by translating ROIs 10-pixel units in both x and y. We then subtracted from the first set of intensity measurements to get net intensity values. After repeated measurements, we imported the net data into MATLAB (The MathWorks) and parsed data values according to thresholds set based on positive and negative controls to determine the intensity distributions and fractions of colocalization.

A caveat of the bead density-based isolation approach is that there is potential for beads that were not internalized or contained within vesicles to have been mixed in with the cell lysate and collected along with the bead-containing vesicles on the sucrose gradient. To gain a more accurate estimate of the average fraction of beads in the 30Q and 81Q isolated populations that were internalized and contained in vesicles, we implemented negative and positive controls. The first of the negative controls consisted of immunofluorescence excluding the primary antibody incubation to assess the validity of the Atto-488 and Alexa-647 secondary fluorescent probes. The fraction of association of the Atto-488 and Alexa-647 probe with 30Q/81Q BDNF-endosomes was 5.00%. The second of the negative controls was a high-salt, 0.25 M MgCl_2_ buffer incubation of the isolated BDNF-endosomes to lower membrane binding-affinity, followed by the immunofluorescence using RAB5/RAB7 primary antibodies. RAB5/RAB7 labels are known to tightly bind membranes and were present in both 30Q and 81Q BDNF-endosome populations^68^. The fraction of association was 8.00% for 30Q endosomes and 4.00% for 81Q endosomes in the MgCl_2_ condition (**Figure S7A, B**). We used intensity distributions and gaussian mixture models to examine trends in the negative control populations to determine a minimum threshold of 300 a.u. for intensity signal to be considered positive. To determine the maximum threshold for association with a single BDNF-endosome, we used a positive control. We employed two-channel immunofluorescence with kinesin-1 and dynein motors, EEA1 and LAMP1, RAB5 and RAB7, RAB11 and RAB7 antibodies for 30Q and 81Q endosome populations. BDNF-beads that associated with either label of interest or a combination of the two at >300 a.u. intensities were considered positive. Based on the average fraction of association taken from the most abundant protein combination, RAB11/RAB7, we estimated that ∼68% of total measurements were from endosomes in the 30Q population, and ∼80% in the 81Q population. We confirmed these estimates with CellTracker Orange (Millipore Sigma) fluorescence colocalization with 560 nm excitation and a minimum threshold of >100 a.u., near the first peak’s mean value using gaussian fits to the intensity distributions. With CellTracker Orange, 68% of total measurements were from endosomes in the 30Q population and 73% in the 81Q population. Intensity distributions and gaussian mixture models for the positive control populations determined a maximum threshold of 5000 a.u. for intensity signal to be considered positive and associated with individual endosomes **(Figure S7A-B).**

### Immunofluorescence Staining Reagent Dilutions

### TrackMate Analysis

#### Parameters

We chose tracking parameters by comparing the results from TrackMate with the maximum intensity projection of the data to ensure that the parameters were optimized to capture the true motility of each cargo. Additionally, we chose parameters that were realistic given the exposure time of the image stack and maximum speed of a motor protein. Given these constraints, we ended up with the parameters presented in **Table 2**.

**Table 2.**
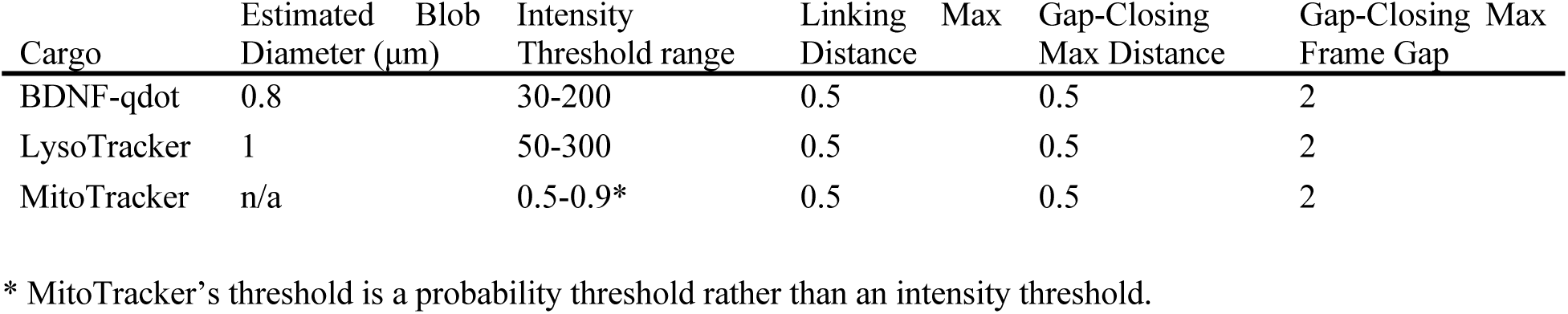
TrackMate parameters according to cargo label.

To account for the irregular shape of mitochondria, we used Weka Segmentation with a custom model to identify and track neuronal mitochondria. To account for photobleaching, we trained the model using the first and last frames of 10 different time-series recordings of neuronal mitochondria. Each mitochondrion was classified as class 1 in the model, while the rest of the axon and the background were identified as class 2. Using this model file, we could use TrackMate with a probability threshold of 0.5-0.9 and the same linking, gap closing, and gap parameters as the other cargoes (**Table 2**) to reliably measure mitochondrial trajectories.

We calculated the processivity parameter α from the slope of the log-log plot of mean squared displacement and averaged the value from all trajectories for each cell. An α=1 indicates purely diffusive behaviour, while active transport is α > 1 and confinement or stationary tracks have α < 1. To determine radius of gyration (R_g_), we used the equation below, where N is the number of points in the trajectory, r_k_ is the current position being evaluated, and r_mean_ is the average position in the trajectory.

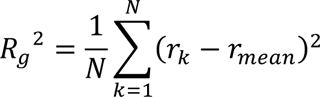

We calculated the tracking uncertainty from TrackMate for each cargo by isolating trajectories from all cargoes that had displacements <0.5 μm and 0 < α > 0.5 and plotting the resultant distribution. We fit a Gaussian mixture model to the distribution and identified the component containing the highest proportion of the data. The mean of the highest proportion component gave the tracking uncertainty for each cargo type.

#### Kymograph Generation

For each kymograph, we manually traced a segmented line along the axon in ImageJ using the segmented line tool with a linewidth of 5. We used this segmented line region of interest to calculate axon segment length. We then used the Multi-Kymograph function on ImageJ to generate the kymograph for the entire image stack (1479 frames). We generated colour-coded kymographs using KymographClear^69^.

#### KymoButler Analysis

We submitted all kymographs to the KymoButler^35^ website with the settings defined in **Table 3** below according to the cargo. We then used the coordinates from each track on each kymograph to generate the trajectories for analysis.

**Table 3.** KymoButler analysis settings.

| Cargo | Sensitivity | Minimum # of frames | µm per pixel | Time int. per frame (s) |
| --- | --- | --- | --- | --- |
| BDNF | 0.1 | 5 | 0.16 | 0.12 |
| Mitochondria | 0.1-0.2 | 5 | 0.16 | 0.12 |
| Lysosome | 0.2 | 10 | 0.16 | 0.12 |

#### Flux Analysis

In MATLAB using the axon segment length determined from the kymograph generation, we divided the segment into fifths for each cell. From each 1/5^th^ position, we identified trajectories from the KymoButler analysis that were located within or passed through the position +/-2.5 μm. We then investigated whether the trajectory started inside the theoretical 5 μm box or outside it. If the trajectory started within the box, we identified it as anterograde flux if it exited from the distal side, retrograde flux if it exited from the somal side, and stationary if it stayed inside the box. If a trajectory entered the box from outside during the imaging time, we identified flux based on its exit location such that those exiting on the distal side were considered anterograde and those exiting on the somal side were considered retrograde. In the case that a trajectory entered from outside of the box and then stayed within it, the flux was identified based on its entry location with anterograde flux for those that entered from the somal side and retrograde flux for those that entered from the distal side. This code generates a binary answer for each condition tested. We added together all anterograde, retrograde, and stationary fluxes from each of the 1/5^th^ position boxes from a given cell and generated an average of these values for each cell. For the multiple box size analysis, we repeated this process varying the box width.

#### Immunofluorescence image analysis

We drew linescans across the 100x images of axons with a linewidth of 10 pixels. We gathered intensity values for tubulin, HTT and <u>DAPI</u> using the measure function in ImageJ. We then calculated the mean intensities for each line profile and the variance in MATLAB.

To quantify neurite arborization length from the 10x images, we first performed a flat-field correction to the image. We then identified neurons by applying an automatic threshold (Otsu) using the Tubb-3 staining. Next, we skeletonized the Tubb-3 channel and measured the percentage of the image area covered by the skeleton. Using the analyze particles function in ImageJ, we determined the number of nuclei from the thresholded DAPI channel. We finally calculated the arborization by dividing the skeleton area by the number of nuclei. To calculate the normalized HTT intensity throughout the neurons, we created a mask of the Tubb-3 channel using gaussian blur and Li automatic threshold functions in ImageJ. We avoided background signal by measuring the HTT and Tubb-3 intensity only within the masked neuron area. Finally, we normalized the HTT fluorescence intensity by dividing the HTT by Tubb-3 values.

#### Stepwise photobleaching analysis

SWPB is an imaging technique that uses the rate of fluorescence signal decay over time to quantify protein complex stoichiometry. We employed immunofluorescence, using a primary target and secondary fluorescent probe to identify proteins of interest. We incubated endosomes in flow chambers for 1 hour and treated them with Pluronic F-127 as described in *Methods: Immunofluorescence imaging*. We used separate chambers for each protein counted. The antibodies we incubated endosomes with included kinesin-1, dynein (monoclonal), HAP1, HTT for 1 hour at RT in the dark. We washed chambers 3X with MAB and incubated with anti-mouse Alexa647 secondary antibody for 1 hour. Following, we washed chambers 3X with MAB. To image the endosomes, we used epifluorescence with 500 ms exposures using a 640 nm laser at 3 mW. We applied a constant laser power applied to each sample for approximately 8 minutes in epifluorescence to induce photobleaching over time. Our single-molecule kinesin-1 control experiment was derived from recent literature in this laboratory^52^. We diluted purified recombinant rat kinesin-1 (rkin430-GFP) to 10 nM and incubated with primary and secondary antibodies and it bound to unlabeled microtubules using 1mM AMPNP to detect single statically bound molecules. Images were acquired using TIRF microscopy.

In FIJI, we selected ROIs with the elliptical selection tool and tightly fitted them to not include any background signal. We compared fluorescent ROIs to brightfield images of beads to verify colocalization and recorded the mean intensity of the fluorescent signal over time. We translated ROIs to an area of local background with no fluorescent signal and measured the mean intensity over time. Next, we subtracted background intensities from their corresponding ROI origin fluorescent signal values and produced net fluorescent signal decay traces. To determine the number and size of discrete bleaching event “steps”, we applied a step-finding algorithm to each trace ^27,28^. We divided the difference between the initial and final fluorescence intensity by the mean step size to get the number of steps. To generate the number of proteins associated with an endosome in the population, we divided the mean step size for a protein of interest by the mean step size for single kinesin-1 **(Figure S7H-I)**. We generated the range of average number steps and proteins per endosome for 30Q and 81Q BDNF-endosome populations using a 95% confidence interval.

#### Estimation of scaffold protein combinations on endosomes

To model the ratio of each protein of interest: dynein, kinesin-1, HAP1 and HTT on BDNF-endosomes, we generated a population of simulated endosomes from our experimental data. From the data in **Figure 7B-D**, stochastic combinations of protein type and number were generated for individual endosomes using random sampling along the cumulative distribution functions of each protein. Bootstrapping was used to determine the mean and 95% confidence intervals of the total number of proteins on each endosome^70^.

### Statistical Analysis

We performed statistical testing for live cell organelle imaging analysis in MATLAB using the one-way ANOVA function and Tukey’s post-hoc test at 95% confidence. We reported p values from the post-hoc tests.

We presented all stepwise photobleaching and fluorescence colocalization data with error bars indicating standard error of the mean (SEM) or 95% confidence intervals as specified in the figure legends. We used bootstrapping analysis in MATLAB to test for statistical significance and determine confidence intervals.

## Author Contributions

E. N. P. P., B. A. T., and A. G. H. designed the study with input from H. M. and G. J. B.. M. A. P. donated cell lines and provided comments on the manuscript. E. N. P. P. and B. A. T. generated neurons through cell inductions. E. N. P. P. performed live cell experiments and analysis. B. A. T. performed BDNF-endosome isolation, experiments, and analysis. M. S. imaged and analyzed fixed neurons at low magnification and provided comments on the manuscript. G. J. B. provided comments on experimental design and the manuscript. D. B. authored in vitro analysis codes and provided input into in vitro experiment design. E. N. P. P., B. A. T., and A. G. H. wrote the manuscript.

## Acknowledgements

1. B. A. T. is supported by a CGS-M fellowship from the National Sciences and Engineering Council of Canada. A. G. H. is supported by the Canadian Institutes of Health Research (PJT-159490 and PJT-185997).

## References

1. Gil, J. M. & Rego, A. C. Mechanisms of neurodegeneration in Huntington’s disease. European Journal of Neuroscience (2008) doi:10.1111/j.1460-9568.2008.06310.x.

2. Ehinger, Y. et al. Huntingtin phosphorylation governs BDNF homeostasis and improves the phenotype of Mecp2 knockout mice . EMBO Mol. Med. (2020) doi:10.15252/emmm.201910889.

3. Her, L. S. & Goldstein, L. S. B. Enhanced sensitivity of striatal neurons to axonal transport defects induced by mutant huntingtin. J. Neurosci. (2008) doi:10.1523/JNEUROSCI.4144-08.2008.

4. Wong, Y. C. & Holzbaur, E. L. F. The regulation of autophagosome dynamics by huntingtin and HAP1 is disrupted by expression of mutant huntingtin, leading to defective cargo degradation. J. Neurosci. (2014) doi:10.1523/JNEUROSCI.1870-13.2014.

5. Caviston, J. P., Ross, J. L., Antony, S. M., Tokito, M. & Holzbaur, E. L. F. Huntingtin facilitates dynein/dynactin-mediated vesicle transport. Proc. Natl. Acad. Sci. U. S. A. 104, 10045–10050 (2007).

6. Engelender, S. Huntingtin-associated protein 1 (HAP1) interacts with the p150Glued subunit of dynactin. Hum. Mol. Genet. (1997) doi:10.1093/hmg/6.13.2205.

7. Li, S. H., Gutekunst, C. A., Hersch, S. M. & Li, X. J. Interaction of Huntingtin-associated protein with dynactin P150(Glued). J. Neurosci. (1998) doi:10.1523/jneurosci.18-04-01261.1998.

8. Colin, E. et al. Huntingtin phosphorylation acts as a molecular switch for anterograde/retrograde transport in neurons. EMBO J. (2008) doi:10.1038/emboj.2008.133.

9. Twelvetrees, A. E. et al. Delivery of GABAARs to Synapses Is Mediated by HAP1-KIF5 and Disrupted by Mutant Huntingtin. Neuron 65, 53–65 (2010).

10. Pal, A., Severin, F., Lommer, B., Shevchenko, A. & Zerial, M. Huntingtin-HAP40 complex is a novel Rab5 effector that regulates early endosome motility and is up-regulated in Huntington’s disease. J. Cell Biol. 172, 605–618 (2006).

11. Seefelder, M., Klein, F. A. C., Landwehrmeyer, B., Fernández-Busnadiego, R. & Kochanek, S. Huntingtin and Its Partner Huntingtin-Associated Protein 40: Structural and Functional Considerations in Health and Disease. J. Huntingtons. Dis. 11, 227–242 (2022).

12. Sahlender, D. A. et al. Optineurin links myosin VI to the Golgi complex and is involved in Golgi organization and exocytosis. J. Cell Biol. (2005) doi:10.1083/jcb.200501162.

13. Evans, C. S. & Holzbaur, E. L. F. Lysosomal degradation of depolarized mitochondria is rate-limiting in OPTN-dependent neuronal mitophagy. Autophagy 16, 962–964 (2020).

14. Gauthier, L. R. et al. Huntingtin controls neurotrophic support and survival of neurons by enhancing BDNF vesicular transport along microtubules. Cell 118, 127–138 (2004).

15. Cason, S. E. et al. Dynein effectors regulate autophagosome motility Sequential dynein effectors regulate axonal autophagosome motility in a maturation-dependent pathway. J. Chem. Biol. (2021).

16. Ayloo, S., Guedes-Dias, P., Ghiretti, A. E. & Holzbaur, E. L. F. Dynein efficiently navigates the dendritic cytoskeleton to drive the retrograde trafficking of BDNF/TrkB signaling endosomes. Mol. Biol. Cell (2017) doi:10.1091/mbc.E17-01-0068.

17. Carabalona, A., Hu, D. J. K. & Vallee, R. B. KIF1A inhibition immortalizes brain stem cells but blocks BDNF-mediated neuronal migration. Nat. Neurosci. (2016) doi:10.1038/nn.4213.

18. Chang, D. T. W., Rintoul, G. L., Pandipati, S. & Reynolds, I. J. Mutant huntingtin aggregates impair mitochondrial movement and trafficking in cortical neurons. Neurobiol. Dis. 22, 388–400 (2006).

19. Orr, A. L. et al. N-terminal mutant huntingtin associates with mitochondria and impairs mitochondrial trafficking. J. Neurosci. 28, 2783–2792 (2008).

20. Song, W. et al. Mutant Huntingtin Binds the Mitochondrial Fission. 17, 377–382 (2011).

21. Trushina, E. et al. Mutant Huntingtin Impairs Axonal Trafficking in Mammalian Neurons In Vivo and In Vitro. Mol. Cell. Biol. 24, 8195–8209 (2004).

22. Caviston, J. P., Zajac, A. L., Tokito, M. & Holzbaur, E. L. F. Huntingtin coordinates the dynein-mediated dynamic positioning of endosomes and lysosomes. Mol. Biol. Cell (2011) doi:10.1091/mbc.E10-03-0233.

23. Ooi, J. et al. Unbiased Profiling of Isogenic Huntington Disease hPSC-Derived CNS and Peripheral Cells Reveals Strong Cell-Type Specificity of CAG Length Effects. Cell Rep. 26, 2494–2508.e7 (2019).

24. Barnat, M. et al. Huntington’s disease alters human neurodevelopment. Science (80-. ). 369, 787–793 (2020).

25. Molina-Calavita, M. et al. Mutant huntingtin affects cortical progenitor cell division and development of the mouse neocortex. J. Neurosci. 34, 10034–10040 (2014).

26. Godin, J. D. et al. Huntingtin Is Required for Mitotic Spindle Orientation and Mammalian Neurogenesis. Neuron 67, 392–406 (2010).

27. Ferrer, I., Goutan, E., Marín, C., Rey, M. J. & Ribalta, T. Brain-derived neurotrophic factor in Huntington disease. Brain Res. 866, 257–261 (2000).

28. Gauthier, L. R. et al. Huntingtin controls neurotrophic support and survival of neurons by enhancing BDNF vesicular transport along microtubules. Cell (2004) doi:10.1016/j.cell.2004.06.018.

29. Wu, L. L. Y., Fan, Y., Li, S., Li, X. J. & Zhou, X. F. Huntingtin-associated protein-1 interacts with pro-brain-derived neurotrophic factor and mediates its transport and release. J. Biol. Chem. 285, 5614–5623 (2010).

30. Yang, M., Lim, Y., Li, X., Zhong, J. H. & Zhou, X. F. Precursor of brain-derived neurotrophic factor (proBDNF) forms a complex with huntingtin-associated protein-1 (HAP1) and sortilin that modulates proBDNF trafficking, degradation, and processing. J. Biol. Chem. 286, 16272–16284 (2011).

31. Yu, C. et al. Decreased BDNF Release in Cortical Neurons of a Knock-in Mouse Model of Huntington’s Disease. Sci. Rep. 8, 1–11 (2018).

32. Baydyuk, M. & Xu, B. BDNF signaling and survival of striatal neurons. Front. Cell. Neurosci. 8, 1–10 (2014).

33. Zhao, X. et al. TRiC subunits enhance BDNF axonal transport and rescue striatal atrophy in Huntington’s disease. Proc. Natl. Acad. Sci. U. S. A. 113, E5655–E5664 (2016).

34. Tinevez, J. Y. et al. TrackMate: An open and extensible platform for single-particle tracking. Methods 115, 80–90 (2017).

35. Jakobs, M. A., Dimitracopoulos, A. & Franze, K. Kymobutler, a deep learning software for automated kymograph analysis. Elife 8, 1–19 (2019).

36. Prowse, E. N. P., Chaudhary, A. R., Sharon, D. & Hendricks, A. G. Huntingtin S421 phosphorylation increases kinesin and dynein engagement on early endosomes and lysosomes. Biophys. J. 122, 1168–1184 (2023).

37. Martinez-Vicente, M. et al. Cargo recognition failure is responsible for inefficient autophagy in Huntington’s disease. Nat. Neurosci. (2010) doi:10.1038/nn.2528.

38. Canty, J. T., Hensley, A., Aslan, M., Jack, A. & Yildiz, A. TRAK adaptors regulate the recruitment and activation of dynein and kinesin in mitochondrial transport. Nat. Commun. 14, 1–15 (2023).

39. Fenton, A. R., Jongens, T. A. & Holzbaur, E. L. F. F. Mitochondrial adaptor TRAK2 activates and functionally links opposing kinesin and dynein motors. Nat. Commun. 12, 4578 (2021).

40. Glater, E. E., Megeath, L. J., Stowers, R. S. & Schwarz, T. L. Axonal transport of mitochondria requires milton to recruit kinesin heavy chain and is light chain independent. J. Cell Biol. 173, 545–557 (2006).

41. van Spronsen, M. et al. TRAK/Milton Motor-Adaptor Proteins Steer Mitochondrial Trafficking to Axons and Dendrites. Neuron 77, 485–502 (2013).

42. Díaz-Hernández, M., Martín-Aparicio, E., Avila, J., Hernández, F. & Lucas, J. J. Enhaced induction of the immunoproteasome by interferon gamma in neurons expressing mutant huntingtin. Neurotox. Res. 6, 463–468 (2004).

43. Nguyen, M., et al. Parkinson ’ s genes orchestrate pyroptosis through selective trafficking of mtDNA to leaky lysosomes. bioRxiv (2023).

44. Strickland, M. R. et al. Ifngr1 and Stat1 mediated canonical Ifn-γ signaling drives nigrostriatal degeneration. Neurobiol. Dis. 110, 133–141 (2018).

45. Warre-Cornish, K. et al. Interferon-γ signaling in human iPSC–derived neurons recapitulates neurodevelopmental disorder phenotypes. Sci. Adv. 6, 1–16 (2020).

46. El-Daher, M., et al. Huntingtin proteolysis releases non-polyQ fragments that cause toxicity through dynamin 1 dysregulation. EMBO J. 34, 2255–2271 (2015).

47. Mutvei, A. P., Nagiec, M. J. & Blenis, J. Balancing lysosome abundance in health and disease. Nat. Cell Biol. 3, (2023).

48. Lim, Y. et al. HAP1 Is Required for Endocytosis and Signalling of BDNF and Its Receptors in Neurons. Mol. Neurobiol. 55, 1815–1830 (2018).

49. Caviston, J. P. & Holzbaur, E. L. F. Huntingtin as an essential integrator of intracellular vesicular trafficking. Trends Cell Biol. 19, 147–155 (2009).

50. Chen, Y., Deffenbaugh, N. C., Anderson, C. T. & Hancock, W. O. Molecular counting by photobleaching in protein complexes with many subunits: Best practices and application to the cellulose synthesis complex. Mol. Biol. Cell 25, 3630–3642 (2014).

51. Chaudhary, A. R., Berger, F., Berger, C. L. & Hendricks, A. G. Tau directs intracellular trafficking by regulating the forces exerted by kinesin and dynein teams. Traffic 19, 111–121 (2018).

52. Beaudet, D. & Hendricks, A. G. Reconstitution of Organelle Transport Along Microtubules In Vitro. Methods in Molecular Biology vols 2623, Chap (2023).

53. Blocker, A. et al. Molecular requirements for bi-directional movement of phagosomes along microtubules. J. Cell Biol. 137, 113–129 (1997).

54. Kruppa, A. J. & Buss, F. Motor proteins at the mitochondria–cytoskeleton interface. J. Cell Sci. 134, (2021).

55. Pu, J., Guardia, C. M., Keren-Kaplan, T. & Bonifacino, J. S. Mechanisms and functions of lysosome positioning. J. Cell Sci. 129, 4329–4339 (2016).

56. Cason, S. E. & Holzbaur, E. L. F. Selective motor activation in organelle transport along axons. Nat. Rev. Mol. Cell Biol. 23, 699–714 (2022).

57. Virlogeux, A. et al. Increasing brain palmitoylation rescues behavior and neuropathology in Huntington disease mice. Sci. Adv. 7, (2021).

58. Lenoir, S. et al. Pridopidine rescues BDNF/TrkB trafficking dynamics and synapse homeostasis in a Huntington disease brain-on-a-chip model. Neurobiol. Dis. 173, (2022).

59. Liot, G. et al. Mutant huntingtin alters retrograde transport of TrkB receptors in striatal dendrites. J. Neurosci. 33, 6298–6309 (2013).

60. Urnavicius, L. et al. Cryo-EM shows how dynactin recruits two dyneins for faster movement. Nature 554, 202–206 (2018).

61. Diefenbach, R. J., Mackay, J. P., Armati, P. J. & Cunningham, A. L. The C-terminal region of the stalk domain of ubiquitous human kinesin heavy chain contains the binding site for kinesin light chain. Biochemistry 37, 16663–16670 (1998).

62. Wu, L. L. Y. & Zhou, X. F. Huntingtin associated protein 1 and its functions. Cell Adhes. Migr. 3, 71–76 (2009).

63. Gunawardena, S. et al. Disruption of axonal transport by loss of huntingtin or expression of pathogenic polyQ proteins in Drosophila. Neuron (2003) doi:10.1016/S0896-6273(03)00594-4.

64. Prigge, J. R. & Schmidt, E. E. HAP1 can sequester a subset of TBP in cytoplasmic inclusions via specific interaction with the conserved TBPCORE. BMC Mol. Biol. 8, 1–18 (2007).

65. Hendricks, A. G., Goldman, Y. E. & Holzbaur, E. L. F. *Reconstituting the motility of isolated intracellular cargoes*. Methods in Enzymology vol. 540 (Elsevier Inc., 2014).

66. Chaudhary, A. R. et al. MAP7 regulates organelle transport by recruiting kinesin-1 to microtubules. J. Biol. Chem. 294, 10160–10171 (2019).

67. Dixit, R. & Ross, J. L. *Studying plus-end tracking at single molecule resolution using TIRF microscopy*. Methods in Cell Biology vol. 95 (Elsevier, 2010).

68. Borchers, A. C., Langemeyer, L. & Ungermann, C. Who’s in control? Principles of rab gtpase activation in endolysosomal membrane trafficking and beyond. J. Cell Biol. 220, 1–15 (2021).

69. Mangeol, P., Prevo, B. & Peterman, E. J. G. KymographClear and KymographDirect: Two tools for the automated quantitative analysis of molecular and cellular dynamics using kymographs. Mol. Biol. Cell 27, 1948–1957 (2016).

70. Beaudet, D., Berger, C. L. & Hendricks, A. G. The sets of kinesins and dynein transporting endocytic cargoes determine the effect of tau on their motility. bioRxiv 2022.06.10.495679 (2023).

