## Supplemental Figures for "Huntingtin polyglutamine expansions misdirect axonal transport by perturbing motor and adaptor recruitment"

### Supplementary Material

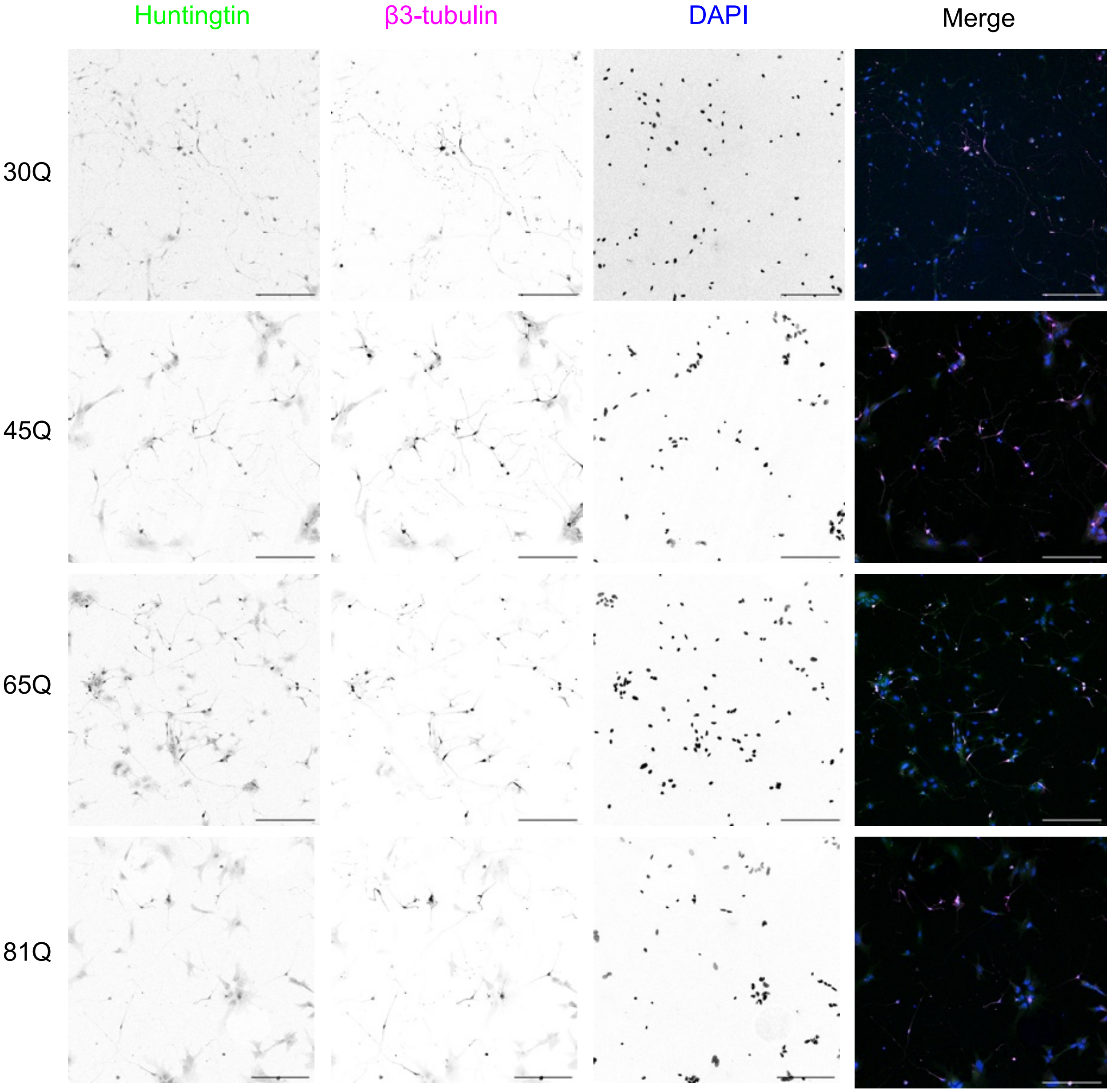


**Figure S1. Immunofluorescence images of HTT, β-3 tubulin, and DAPI at scale.** Images at scale during acquisition used in Figure 1A-C. Cell line is indicated on the left of the row of representative images, fluorophores are indicated above the column of the single channel inverted images and the text is in the colour they are shown in the merged image. Scale bars 200 μm.

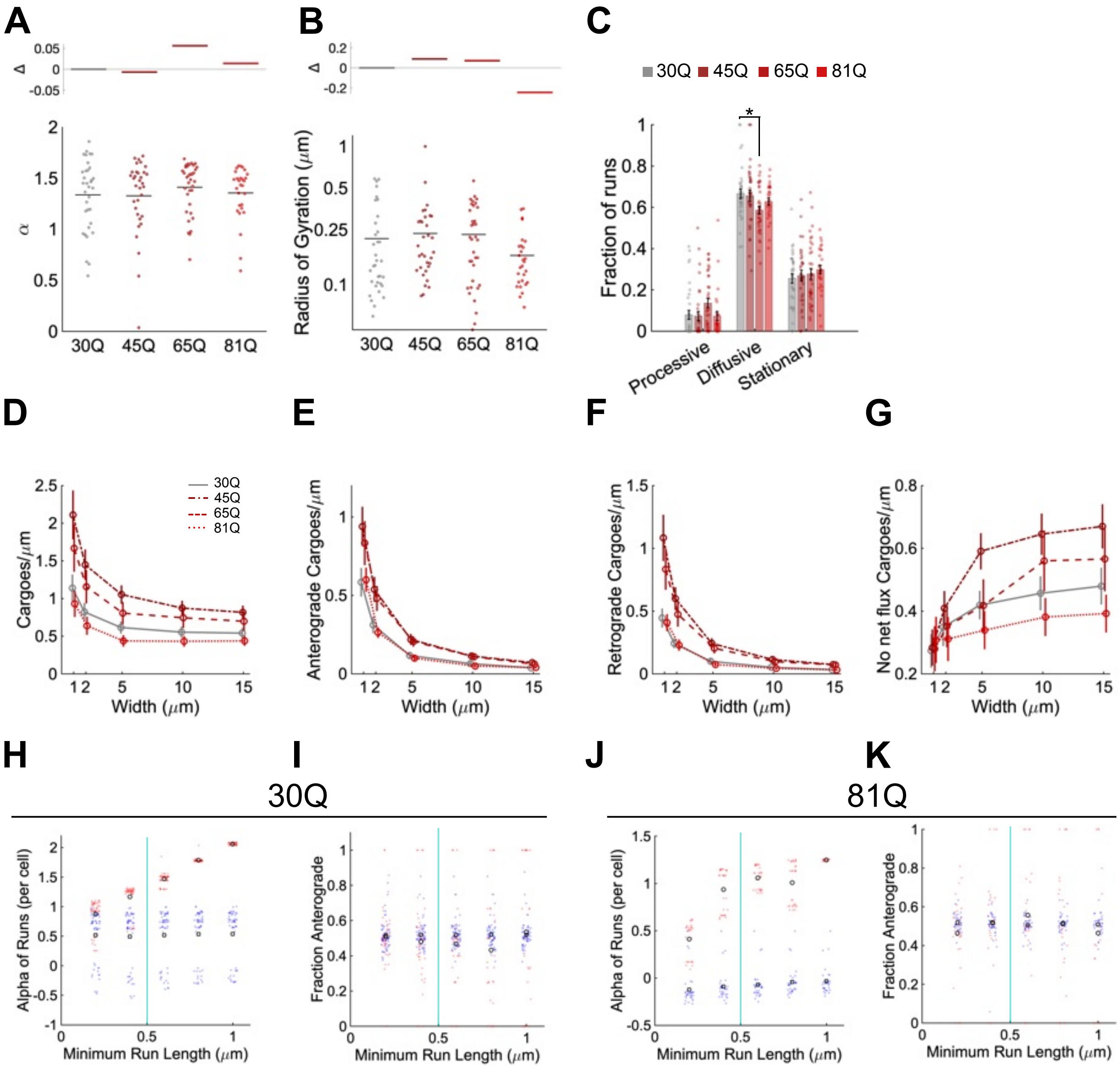


**Figure S2. BDNF cargo short-range transport remains unaffected by HTT polyglutamine length. A-C** Short-range track analysis using TrackMate to identify trajectories. **A** Mean processivity per cell (α) of BDNF vesicles for each cell line, with the fraction change (Δ) indicated by the coloured lines above. **B** Mean displacement per cell by radius of gyration presented on a logarithmic scale, Δ indicated above as in **A**. **C** Mean fraction of processive, diffusive, and stationary runs per cell determined using a minimum processive run length of 0.5 μm, with the colour code above. **D-G** Determining optimal box size for flux analysis. **D** Total number of trajectories that enter the 5 boxes set at the given width, as a measure of the approximate number of cargoes per area in each cell line. Error bars represent the SEM, points indicate the mean. **E-G** represent the number of cargoes from **D** with a net anterograde, retrograde, without flux at each given box width. **H** Processivity per cell (α) of HTT-30Q cell BDNF vesicles with minimum run lengths of 0.2-1 μm, with processive motion indicated in red and mean by a circle, and diffusive motion shown in blue with a square representing the mean. A cyan line indicates the minimum run length chosen for the analysis in **A-C**. **I** Fraction of cargoes moving anterograde compared to retrograde depending on minimum run length threshold, with the same colour code as in **H**. **J,K a**s in **H,I** however for HTT-81Q cells. Statistical significance with $p<0.05$ is indicated by the p value itself or an * and was determined by one-way analysis of variance and a Tukey multiple comparison post-hoc test. The value for the asterisk is $p=0.044$.

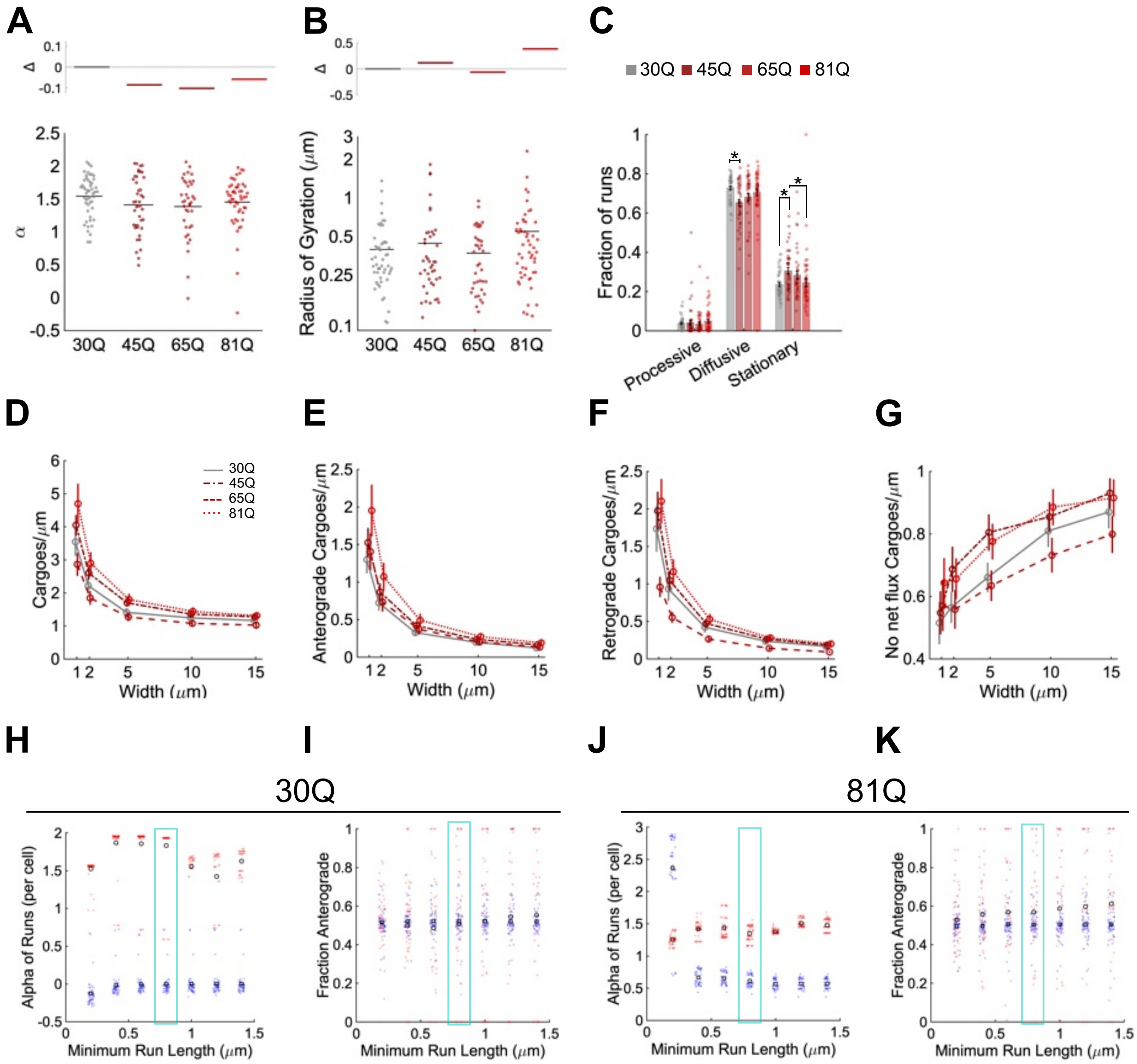


**Figure S3. Lysosome diffusive fraction slightly decreases and stationary fraction increases with polyglutamine HTT. A-C** Short-range track analysis using TrackMate to identify trajectories. **A** Mean processivity per cell (α) of lysosomes for each cell line, with the fraction change (Δ) indicated by the coloured lines above. **B** Mean displacement per cell by radius of gyration presented on a logarithmic scale, Δ indicated above as in **A**. **C** Mean fraction of processive, diffusive, and stationary runs per cell determined using a minimum processive run length of 0.8 µm, with the colour code above. **D-G** Determining optimal box size for flux analysis. **D** Total number of trajectories that enter the 5 boxes set at the given width, as a measure of the approximate number of cargoes per area in each cell line. Error bars represent the SEM, points indicate the mean. **E-G** represent the number of cargoes from **D** with a net anterograde, retrograde, without flux at each given box width. **H** Processivity per cell (α) of HTT-30Q cell lysosomes with minimum run lengths of 0.2-1.4 µm, with processive motion indicated in red and mean by a circle, and diffusive motion shown in blue with a square representing the mean. A cyan box indicates the minimum run length chosen for the analysis in **A-C**. **I** Fraction of cargoes moving anterograde compared to retrograde depending on minimum run length threshold, with the same colour code as in **H**. **J,K a**s in **H,I** however for HTT-81Q cells. Statistical significance with $p<0.05$ is indicated by the p value itself or an * and was determined by one-way analysis of variance and a Tukey multiple comparison post-hoc test. The values for those with asterisks are for diffusive fraction 30Q vs 45Q p=0.0064 and for stationary fraction 30Q vs 45Q p=0.016, stationary fraction 45Q vs 81Q p=0.045.

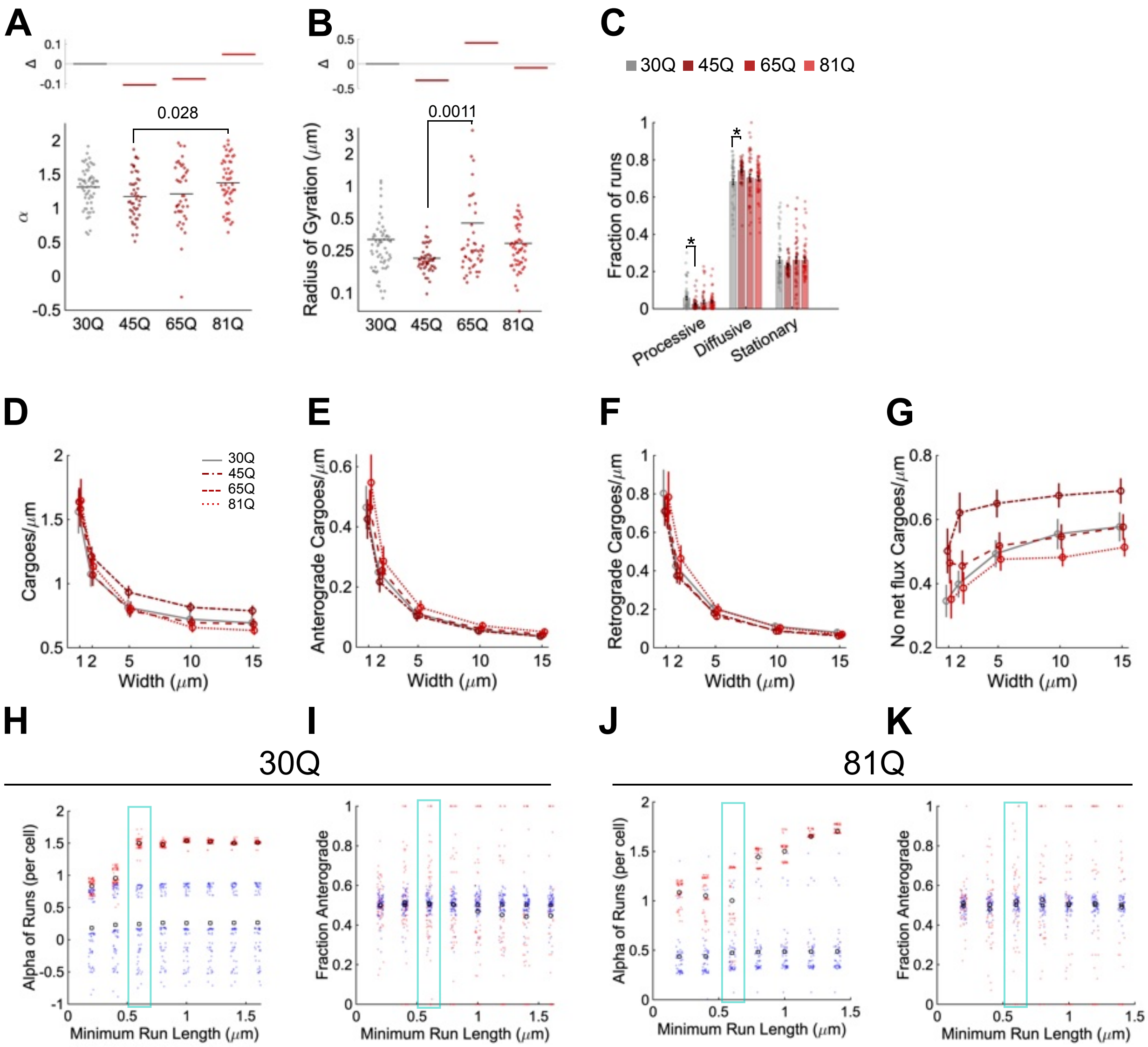

**Figure S4. Mitochondrion transport is slightly less processive with pathogenic HTT. A-C** Short-range track analysis using TrackMate to identify trajectories. **A** Mean processivity per cell (α) of mitochondria for each cell line, with the fraction change (Δ) indicated by the coloured lines above. **B** Mean displacement per cell by radius of gyration presented on a logarithmic scale, Δ indicated above as in **A**. **C** Mean fraction of processive, diffusive, and stationary runs per cell determined using a minimum processive run length of 0.5 µm, with the colour code above. **D-G** Determining optimal box size for flux analysis. **D** Total number of trajectories that enter the 5 boxes set at the given width, as a measure of the approximate number of cargoes per area in each cell line. Error bars represent the SEM, points indicate the mean. **E-G** represent the number of cargoes from **D** with a net anterograde, retrograde, without flux at each given box width. **H** Processivity per cell (α) of HTT-30Q cell mitochondria with minimum run lengths of 0.2-1.6 µm, with processive motion indicated in red and mean by a circle, and diffusive motion shown in blue with a square representing the mean. A cyan box indicates the minimum run length chosen for the analysis in **A-C**. **I** Fraction of cargoes moving anterograde compared to retrograde depending on minimum run length threshold, with the same colour code as in **H**. **J,K** as in **H,I** however for HTT-81Q cells. Statistical significance with $p<0.05$ is indicated by the p value itself or an * and was determined by one-way analysis of variance and a Tukey multiple comparison post-hoc test. The values for those with asterisks are for processive motility 30Q vs 45Q $p=0.013$, and for diffusive motility 30Q vs 45Q $p=0.014$.

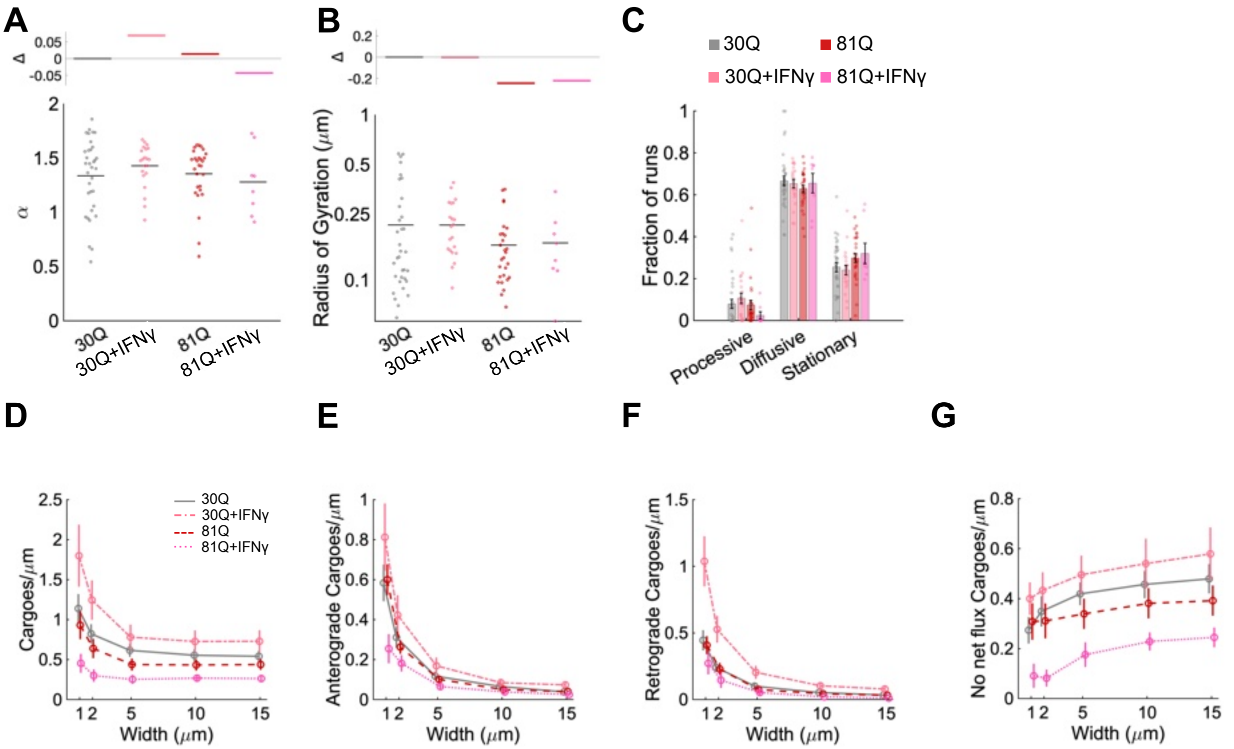


**Figure S5. Stress mildly decreases BDNF processivity and displacement with pathogenic HTT-polyQ.** Data from the same images as in Figure 5: 30Q (same samples as Figure 2, n=56, 4 terminal differentiations), 30Q with 24 hrs 100 ng/mL IFγ (n=20, 3 terminal differentiations), 81Q (same samples as Figure 2, n=34, 4 terminal differentiations), and 81Q with 24 hrs 100 ng/mL IFγ (n=15, 3 terminal differentiations). **A-C** Short-range track analysis using TrackMate to identify trajectories. **A** Mean processivity per cell (α) of BDNF vesicles for each condition, with the fraction change (Δ) indicated by the coloured lines above. **B** Mean displacement per cell by radius of gyration presented on a logarithmic scale, Δ indicated above as in **A**. **C** Mean fraction of processive, diffusive, and stationary runs per cell determined using a minimum processive run length of 0.5 μm, with the colour code above. **D-G** Determining optimal box size for flux analysis. **D** Total number of trajectories that enter the 5 boxes set at the given width, as a measure of the approximate number of cargoes per area in each cell line. Error bars represent the SEM, points indicate the mean. **E-G** represent the number of cargoes from **D** with a net anterograde, retrograde, without flux at each given box width. Statistical significance with $p<0.05$ is indicated by the p value itself and was determined by one-way analysis of variance and a Tukey multiple comparison post-hoc test.

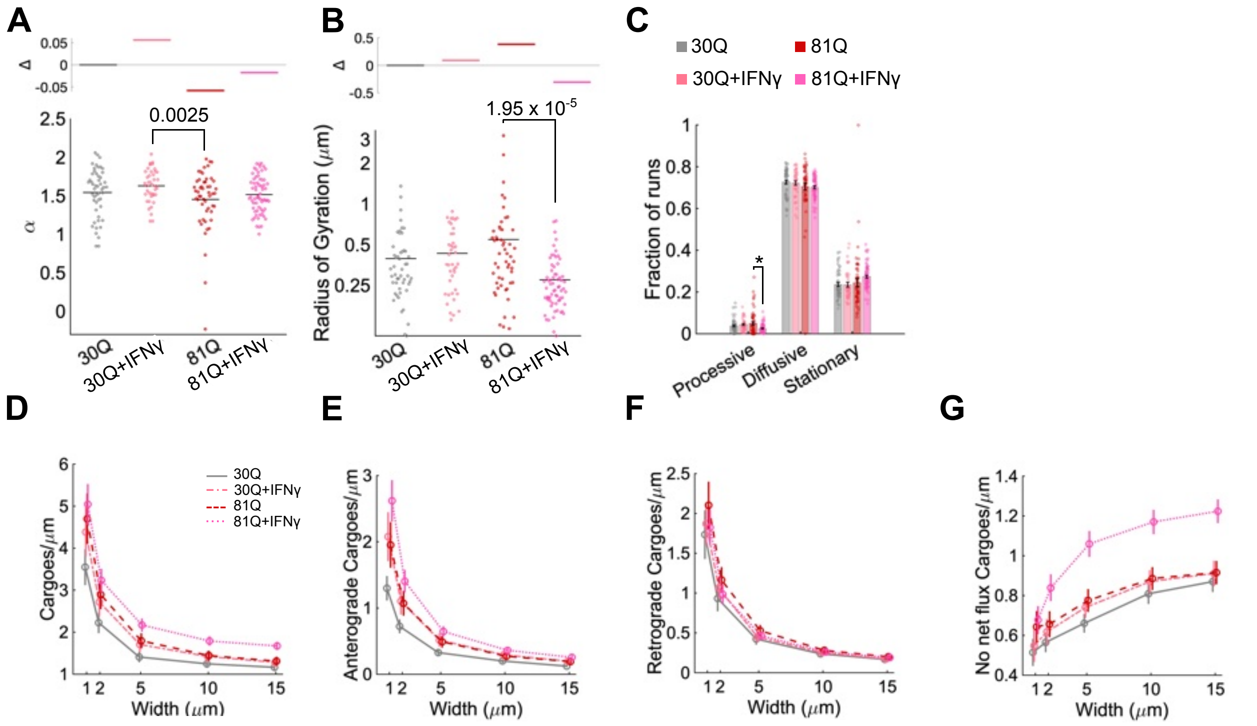


**Figure S6. Stress decreases displacement and decreases the fraction of lysosomes that are processive only in cells containing pathogenic HTT. A-C** Short-range track analysis using TrackMate to identify trajectories. **A** Mean processivity per cell (α) of BDNF vesicles for each cell line, with the fraction change (Δ) indicated by the coloured lines above. **B** Mean displacement per cell by radius of gyration presented on a logarithmic scale, Δ indicated above as in **A**. **C** Mean fraction of processive, diffusive, and stationary runs per cell determined using a minimum processive run length of 0.8 μm, with the colour code above. **D-G** Determining optimal box size for flux analysis. **D** Total number of trajectories that enter the 5 boxes set at the given width, as a measure of the approximate number of cargoes per area in each cell line. Error bars represent the SEM, points indicate the mean. **E-G** represent the number of cargoes from **D** with a net anterograde, retrograde, without flux at each given box width. Statistical significance with $p<0.05$ is indicated by the p value itself or an * and was determined by one-way analysis of variance and a Tukey multiple comparison post-hoc test. The value with the asterisk for processive motility 81Q vs 81Q+IFγ p=0.0056.

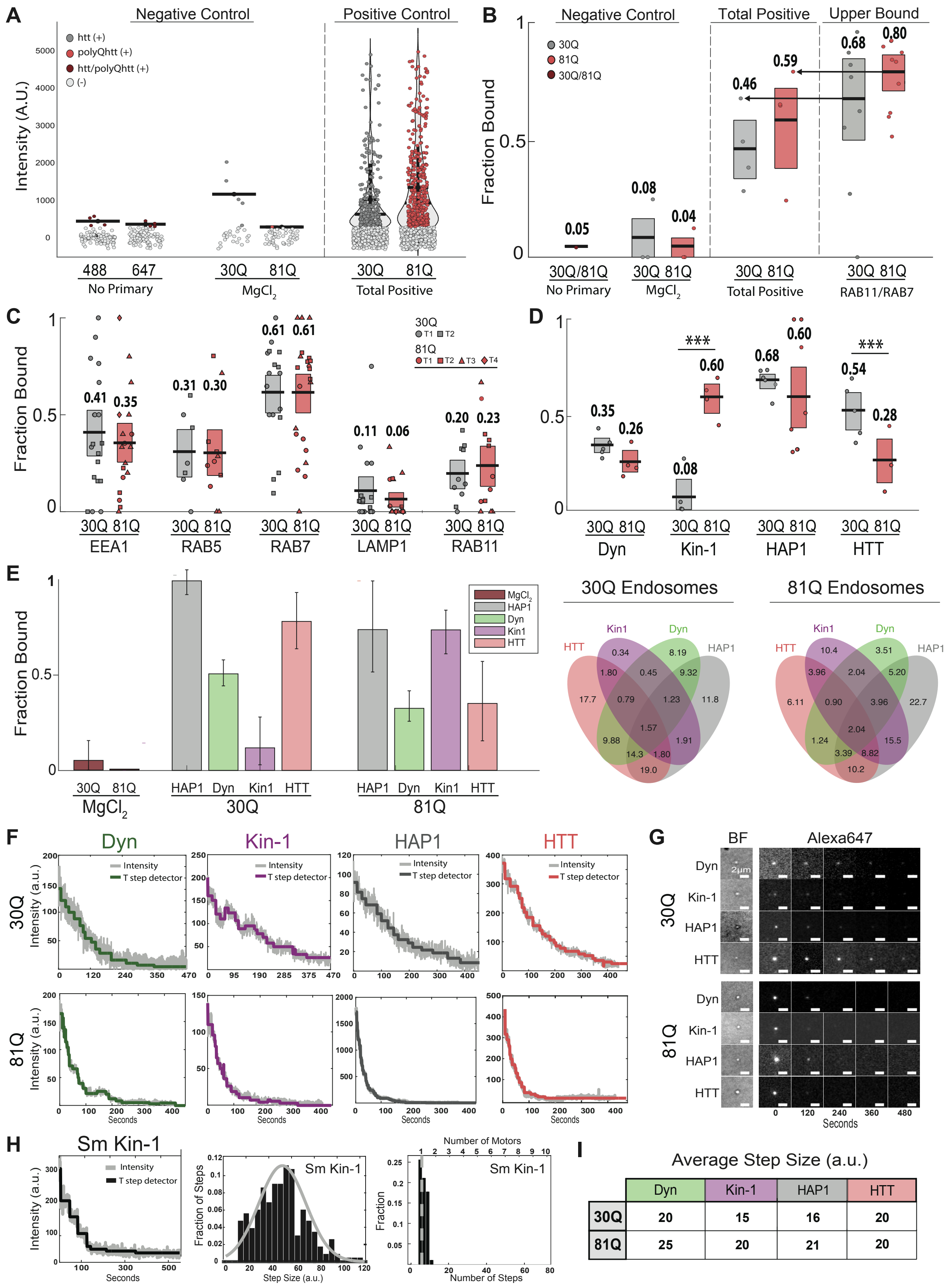


| | Dyn | Kin-1 | HAP1 | HTT |
|---|---|---|---|---|
| 30Q | 20 | 15 | 16 | 20 |
| 81Q | 25 | 20 | 21 | 20 |

**Figure S7. Controls and additional data supporting in vitro assays.**

**A** Intensity distributions for immunofluorescence of BDNF-endosome controls. Fraction of endosomes positive for label of interest for 30Q (grey), positive for 81Q (red), positive for 30Q/81Q (burgundy), and negative for label of interest (light grey). Negative controls consist of a no-primary condition, Atto-488 (n=92) and Alexa-647 (n=89), and a high salt, MgCl2 buffer used to lower vesicle membrane binding affinity. 2-channel positive colocalization controls were generated from motor and membrane-associated proteins (dynein/kinesin-1, EEA1/LAMP1, RAB5/RAB7, RAB11/RAB7) (see methods). **B** Fractions of colocalization for 2-channel negative and positive controls. No primary condition showed 5.00% secondary antibody fraction of association with 30Q/81Q BDNF-endosomes. MgCl2 condition showed 8.00% fraction of association in 30Q (n=27) and 4.00% fraction of association in 81Q (n=50) BDNF-endosomes, with each data point representing a different field of view. Total positive controls showed 43.0% fraction of association in 30Q (n=628) and 51.9% fraction of association in 81Q (n=728) BDNF-endosomes, with each data point representing a different combination of labels (Figure S7A). RAB11/RAB7 control population exhibited the greatest fraction of colocalization in the total control population 68.0% in 30Q and 80.0% in 81Q, with each datapoint representing a different field of view. Arrows represent the means of RAB11/RAB7 colocalization fraction. **C** Plot shows fraction of isolated BDNF-endosomes positive for dynein (30Q, n=316 endosomes, 5 fields of view, 2 isolations; 81Q, n=162, 4 fields of view, 2 isolations), kinesin-1 (30Q, n=316, 5 fields of view, 2 isolations; 81Q, n=162, 4 fields of view, 2 isolations), HAP1 (30Q, n=498, 6 fields of view, 2 isolations; 81Q, n=142, 8 fields of view, 2 isolations) and HTT (30Q, n=369, 5 fields of view, 2 isolations; 81Q, n=97, 3 fields of view, 1 isolation) for 30Q (dark) and 81Q (light) neurons. **D** Plot shows fraction of isolated BDNF-endosomes positive for membrane-associated protein stains EEA1, RAB5, RAB7, LAMP1 and RAB11 for 30Q (grey) and 81Q (red) neurons corresponding to intensity plots. Black lines represent the mean values and boxes show 95% confidence interval for means generated by bootstrapping with replacement. Shapes represent fraction of positive BDNF-endosomes for each field of view grouped by neuronal terminal differentiation. MANOVA and Independent samples T-test were performed to assess statistical significance ($p < 0.05$). **E** On the right, plot shows the bootstrapped, computational model fractions of isolated 30Q and 81Q BDNF-endosomes positive for dynein (green), kinesin-1 (purple), HAP1 (grey) and HTT (red), adjusted for the respective fraction of bead containing endosomes (30Q=68%, 81Q=80%) (see methods). On the left, Venn diagrams display the ratios of dynein, kinesin 1, HAP1 and HTT exhibiting association with endosomes in each population. **F** SWPB traces generated from intensity (a.u.) decay over time (seconds) are used to estimate step sizes for proteins of interest. Representative photobleaching step-traces of dynein (blue), kinesin-1 (red), HAP1 (green) and HTT (purple) for 30Q (top) and 81Q (bottom) BDNF-endosomes. **G** Brightfield (BF) BDNF-bead and immunofluorescence channel (647 nm) sample frames are shown for each target protein from the HTT scaffold, dynein, kinesin-1, HAP1 and HTT from 0-480 seconds during photobleaching. Scale bars 2 μm. **H** Plots display example SWPB trace generated from intensity (a.u.) decay over time (seconds), step size distribution and average number of steps distribution for single molecule kinesin-1 control (average step size = 50.6 (a.u.); average number of steps = 6.90). **I** Average step size determined from stepwise photobleaching of dynein, kinesin-1, HAP1 and HTT associated with 30Q and 81Q endosomes.
